# An amygdala circuit mediates experience-dependent momentary exploratory arrests

**DOI:** 10.1101/797001

**Authors:** Paolo Botta, Akira Fushiki, Ana Mafalda Vicente, Luke A. Hammond, Alice C. Mosberger, Charles R. Gerfen, Darcy Peterka, Rui M. Costa

## Abstract

Exploration of novel environments ensures survival and evolutionary fitness. This behavior is expressed through exploratory bouts and arrests, which change dynamically based on experience. Neural circuits mediating exploratory behavior should therefore integrate experience and use it to select the proper behavioral output. Using a spatial exploration assay, we uncovered an experience-dependent increase of momentary arrests in visited locations where animals previously arrested. Quantitative analyses of neuronal calcium activity in freely-exploring mice revealed that a large neuronal ensemble in basolateral amygdala is active during self-paced behavioral arrests. This ensemble was recruited in an experience-dependent manner, and closed-loop optogenetic manipulation of these neurons revealed that they are sufficient and necessary to drive experience-dependent arrests. Additionally, we found that neurons in the basolateral amygdala projecting to central amygdala mediate these momentary arrests. These findings uncover an amygdala circuit that mediates momentary exploratory arrests in familiar places, without changing place preference or anxiety/fear-like behaviors.

## Introduction

Exploration is highly conserved across species, and can be construed as the act of gathering information or resources from unknown surroundings (Fonio, Benjamini and Golani, 2009). It is therefore a self-paced process in which animals guide their behavior to learn about new environments. This behavior is fundamental in discovering new territories in order to gather food, water or find a safe shelter, e.g. birds explore the territory before choosing a place to nest (Mills, 1985; Freire, Appleby and Hughes, 1996). Novelty is one of the main drives for exploration, and can override other drives (Fonio, Benjamini and Golani, 2009).

Many past behavioral studies analyzing exploration in the lab used assays that depend on water or food-deprived animals executing movements in order to ultimately find and consume food or water (Fonio, Benjamini and Golani, 2009; Benjamini *et al.*, 2010; Vorhees and Williams, 2014). Less is known about the self-paced moment-to-moment actions that govern the exploration of novel environments in the absence of explicit reinforcers (Tolman, 1948; Renner, 1990; Benjamini *et al.*, 2011; Redish, 2016). Emergence exploratory paradigms were developed in order to study naturalistic exploratory behavior during transitions between familiar and novel environments (Fonio, Benjamini and Golani, 2009; Benjamini *et al.*, 2011). These paradigms allow animals to explore a novel area in a self-paced manner, departing from a familiar one, and permit the investigation of moment-to-moment exploratory dynamics during spatial familiarization as the animal performs defined trips from the familiar shelter. When facing novel surroundings and given the freedom of movement, animals execute gradual and structured exploratory trips with the development of a quantifiable behavioral gradient (Benjamini *et al.*, 2011).

Several reports show that a wide variety of species use specific locations, sometimes named home bases, as strategic points of reference from which the animal begins and terminates exploratory trips (Eilam and Golani, 1989; Golani, Benjamini and Eilam, 1993; Drai, Benjamini and Golani, 2000; Clark, Hamilton and Whishaw, 2006; Fonio, Benjamini and Golani, 2009; Benjamini *et al.*, 2011). In exploration, home bases can be perceived as the origin of exploratory activity at the interface between known and unknown places, therefore representing a key factor in understanding its general structure and organization (Eilam and Golani, 1989; Golani, Benjamini and Eilam, 1993; Clark, Hamilton and Whishaw, 2006; Dvorkin, Szechtman and Golani, 2010). These locations are characterized by more frequent behavioral arrests than other areas explored by an animal (Eilam and Golani, 1989; Golani, Benjamini and Eilam, 1993), indicating that exploratory motor dynamics are influenced by spatial knowledge. Exploratory arrests are distinct from freezing responses, as they are momentary, occur voluntarily in familiar places, and are not triggered by any apparent aversive stimuli (LeDoux and Phillips, 1992; Roelofs, 2017; Roseberry and Kreitzer, 2017). Instead, these type of arrests may be critical for the deliberation process in familiar/safe places, while the animal prepares for the next exploratory trip (Eilam and Golani, 1989; Clark, Hamilton and Whishaw, 2006; Redish, 2016).

Given that behavioral exploratory actions such as arrests depend upon knowledge of the environment, the neural circuits mediating this behavior should integrate experience-dependent contextual information and use it to select a proper behavioral output. The main input structure of the amygdala, the basolateral nucleus (BLA), has been shown to be involved in the learning of contextual information related to appetitive and aversive experiences (LeDoux and Phillips, 1992; Herry *et al.*, 2008; Janak and Tye, 2015; Namburi *et al.*, 2015; Beyeler, Namburi, Gordon F. Glober, *et al.*, 2016; Kim *et al.*, 2016; Behaviors *et al.*, 2017; Beyeler, C. J. Chang, *et al.*, 2018), as well as in the expression of motor defensive behaviors via the central amygdala (CEA, Ciocchi *et al.*, 2010; Botta *et al.*, 2015; Tovote *et al.*, 2016; Xu, Krabbe, Gründemann, *et al.*, 2016; Fadok *et al.*, 2017; Terburg *et al.*, 2018). Previous studies suggest that the amygdala plays a much broader role in integrating novel and familiar sensory stimuli in humans and monkeys; lesions of the BLA impair familiarity recognition (Wilson and Rolls, 1993; Schwartz *et al.*, 2003; Mason *et al.*, 2006; Farovik *et al.*, 2011), and affect locomotor exploration in the open field (Jellestad and Bakke, 1985; Yim and Mogenson, 1989).

In the present study, we developed an emergent assay to investigate the dynamics of exploration in mice, and in particular the neural mechanisms underlying momentary exploratory arrests. Using behavioral analyses, circuit mapping, single-cell calcium imaging and closed-loop optogenetic approaches, we provide evidence for the recruitment of a well-defined BLA neuronal population that mediates experience-dependent momentary arrests. Furthermore, activation and inhibition of the BLA neuronal population directly projecting to CEA revealed that this projection mediates behavioral arrests. The role of amygdala in modulating exploratory arrests was not directly associated with changes in the place value or place preference, nor anxiety/fear-like behavior.

## Results

### A BLA neuronal ensemble is active during self-paced behavioral arrests

We investigated the progression of self-paced exploratory dynamics by analyzing the behavior of animals as they explored a 100cm circular arena, departing from a familiar 20cm-arena, for 5 consecutive days. Every animal was habituated to the small 20cm-arena for 30min (7 mice). Subsequently, to give the animal the chance to explore the 1m-arena without human interference, an automatic gate was opened and left open for 20min (Figure 1A and Figure S1). The arenas were cleaned with 70% ethanol and an odor killer solution at the end of each session (see Star Methods). While animals performed a constant number of trips from the small to the big arena with similar daily preference for the 100cm-arena, the duration of the trips and the area explored gradually increased during the five experimental days (Figure 1B and Figure S1; Movie S1-S2). As shown by our results and others (Benjamini *et al.*, 2011), the complexity of the trajectories increased in late exploration (Figure S1). Our data confirm that the developed behavioral assay can be used to study the gradual progression of exploration.

**Figure 1.**
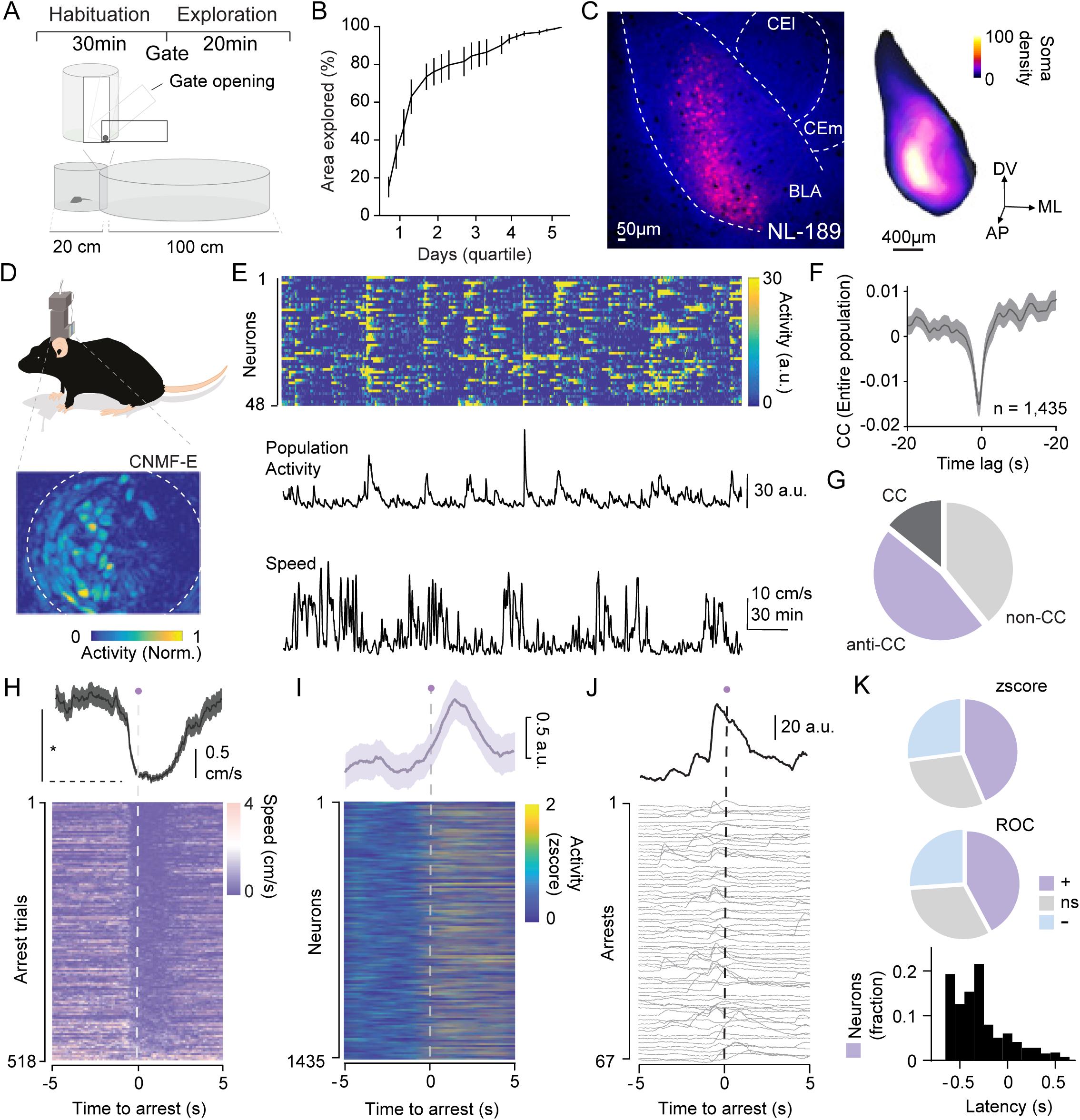
A BLA neuronal ensemble encodes self-paced behavioral arrests. A, Top, behavioral protocol. Bottom, dimensionality emergence assay. B, Daily area explored (%) divided in quartiles, considering even portions of exploratory time. C, Left, brain section showing the unilateral AAV1-CAG-FLEX-Tdtomato injection in BLA of NL189-cre mouse line. Lines denoted the injection site and amygdala borders. Central amygdala, CEA, is shown divided in the lateral, l, and medial, m, part. Right, Heatmap of BLA soma density defined as the number of cells within a 300 micron diameter sphere. D, Top, scheme of mini-endoscope mounted on top of the mouse’s head. Bottom, CNMF-E processed movie of the neurons located in BLA, normalized neuronal calcium activity by the max. E, Top, time course of the activity of many single neurons simultaneously recorded in one animal during the exploration of the large arena. Center, average population activity from the same animal. Bottom, movement speed during exploration. F, Time course of the cross-correlation (CC) for the entire neuronal population at different lags (range -20 to 20s). G, Pie chart showing the proportion of the CC, anti-CC and non-CC neurons (for all neurons/all animals). H, Top, average speed of the arrest trials. Bottom, time course of the speed in all arrest trials for all the animals. Time to arrest (0s) is indicated by a dashed gray line and the purple dot. I, Top, average raw calcium trace of the entire neuronal ^NL189^BLA population imaged (n = 1435) at the time of arrest (in arbitrary unit, a.u.). Bottom, z-score of all ^NL189^BLA single neuron activity at the time of arrest. J, Representative trace of a ROC-sorted arrest neuron showing the average (top, black line) and the single arrest trials (bottom, gray). K, Top, pie chart of z-scored and ROC-sorted neurons; positively (+), negatively (-) and non-modulated (ns). Bottom, Time course of the peak neural activity vs time to arrest. Latency 0s represents the time to arrest. Data are presented as mean ± standard error of the mean (SEM).

In order to precisely target excitatory pyramidal neurons in BLA, we used a mouse line that expresses cre recombinase in a population of BLA neurons - NL189 line (Figure 1C and Figure S2). We targeted NL189 cre neurons (^NL189^BLA neurons) with an adeno-associated virus (AAV) expressing tdTomato in a cre dependent manner to quantify their location in BLA (Figure 1C; Movie S3). Anatomical analysis and automated quantification of cell populations reveal that ^NL189^BLA neurons are primarily located in the lateral-posterior part of the basal amygdala and ventral-lateral part of the lateral amygdala (Figure 1C and Figure S2; Movie S4). Double injection of AAV viruses, one expressing tdTomato in a cre dependent manner and one expressing yellow fluorescence protein (YFP) in a CamKII-dependent manner, showed an overlap of ∼80% between ^NL189^BLA neurons and CamKII-expressing neurons (Figure S3). Using a clustering analysis based on three parameters collected with the input-output function of ^NL189^BLA neurons (first inter-spike interval distribution, spike adaptation and maximum firing rate), we estimated that the majority are regular firing (Figure S3). Together these results confirm the nature of ^NL189^BLA neurons as principal cells in BLA (pyramidal; Faber, Callister and Sah, 2001; Likhtik *et al.*, 2006; Sosulina *et al.*, 2006; Fonio, Benjamini and Golani, 2009; Ehrlich, Ryan and Rainnie, 2012).

We used the adapted emergence behavioral assay to track the moment-to-moment exploratory path of NL189 cre^+^ animals and calculated their movement speed based on *xy* coordinates (Figure S1). After injecting a conditional AAV virus expressing GCamp6f in BLA (Chen *et al.*, 2013), we implanted a gradient index lens in seven mice (Figure S4). Neuronal calcium activity was imaged from 1,435 ^NL189^BLA neurons for five days (average neurons/animal/day = 41 ± 4.85, Figure S4) using a miniaturized one-photon fluorescence microscope in freely-exploring animals and processed with the CNMF-E algorithm (Figure 1D; Figure S4; Movie S5; see Star Methods; Pnevmatikakis *et al.*, 2016). We observed that a large population of simultaneously recorded ^NL189^BLA neurons had coincident, transient moments of increase in activity, which corresponded to moments when the speed of the animals decreased (Figure 1E). In order to understand whether the neuronal activity was correlated with general movement speed during exploration, we compared the z-scored population average activity with the z-scored speed using a cross-correlation analysis previously used (Costa *et al.*, 2006). Overall, the activity of the entire neuronal population of ^NL189^BLA neurons was inversely cross-correlated with speed (Figure 1F). A rather large proportion of single neurons was activated at low speed (anti-CC) compared to high speed (positive cross-correlated) neurons (Figure 1G). Another analytical approach, the spike-triggering average, confirmed a large proportion of ^NL189^BLA neurons active during decreased movement speed (Figure S6). Using a segmentation algorithm (Benjamini *et al.*, 2010; Figure S1; see Star Methods), we were able to detect the transition between movements and behavioral arrests (Figure 1H). These arrests were momentary (average duration in big arena 1.11 ± 0.21 s), and were usually followed by an increase in the angular speed of head, (Figure S5; Movie S6, S7), suggestive of *vicarious trial and error* behavior (Muenzinger and Gentry, 1931; Tolman, 1939; Redish, 2016). In combination with the ROC and z-score analysis to sort neuronal activity between behavioral states, we found that a large proportion of ^NL189^BLA neurons was positively modulated (active) during the transition between movement and momentary arrests compared to negatively-modulated (inhibited) neuronal population (Figure 1I-K). The majority of arrest ^NL189^BLA neurons was active before arrest (latency of -0.29 ± 0.01 s, Figure 1K). Overall, these results show that a large subpopulation of BLA principal neurons, the ^NL189^BLA neurons, is active before and during behavioral arrests.

### Transient ^NL189^BLA neuron activation decreases movement speed and triggers arrest

In order to test if the activity of ^NL189^BLA neurons is causally related to behavioral arrests, we injected a conditional AAV virus expressing channelrhodopsin hChR2(H134R) (ChR) and YFP in BLA. Optogenetic stimulation of ^NL189^BLA neurons using three different light patterns (1x, one 10ms pulse; 2x, two 10ms pulses at 20Hz; 5x, five 10ms pulses at 20Hz; Methods) showed a strong and reliable effect on eliciting neuronal activity in slice (Figure 2A-B and Figure S7). Next, after bilateral injection of the ChR-(ChR group) or only YFP-expressing (Control group) virus in BLA, optical fibers were implanted 200μm above BLA to perform a closed-loop optogenetic stimulation in which the LED_on_ was triggered ∼33% of the time during bouts where the animal was moving and had accelerated for at least 100ms (trial on; Figure 2C-F; see Star Methods). The 66% of trials when light was not delivered served as within-animal controls (off trials; Figure 2F-J). Acceleration-locked closed-loop optogenetic activation of ^NL189^BLA neurons using the three aforementioned optogenetic protocols decreased the speed of the animals tested in comparison to off trials and Control animals (Figure 2F-K and Figure S8; Movie S8). The same stimulation protocol was ineffective in eliciting any evident change in speed in Control animals (Figure 2G-K and Figure S8). Closed-loop optogenetic activation of ^NL189^BLA neurons in the ChR group elicited a significant increase in behavioral arrests following stimulation (Figure 2M). The average duration of the arrests resulting from the 1x, 2x and 5x stimulation protocols was 0.57 ± 0.08 s, 0.64 ± 0.11 s and 0.47 ± 0.08 s respectively, suggesting that these are momentary arrests and not freezing (more prolonged state; (Herry *et al.*, 2008; Tovote *et al.*, 2016). Furthermore, there was no difference between the overall count of arrests throughout the behavioral session for the ChR vs. Control group (Figure 2N), from which it can be deducted that the arrests were not caused by the animal freezing to the environment. Additionally, classical analysis used to detect anxiety-like behavior showed that the most intense optogenetic stimulation (5x) did not have a consistent effect on speed found at different times from LED_on_ stimulation, or distance from the center of the arena (Figure 2L and Figure S8). In summary, direct optogenetic stimulation of ^NL189^BLA neurons during ongoing movement reduces speed and promotes arrests but does not have any impact on overall light-unlocked speed, arrest count or anxiety-like behavior.

**Figure 2.**
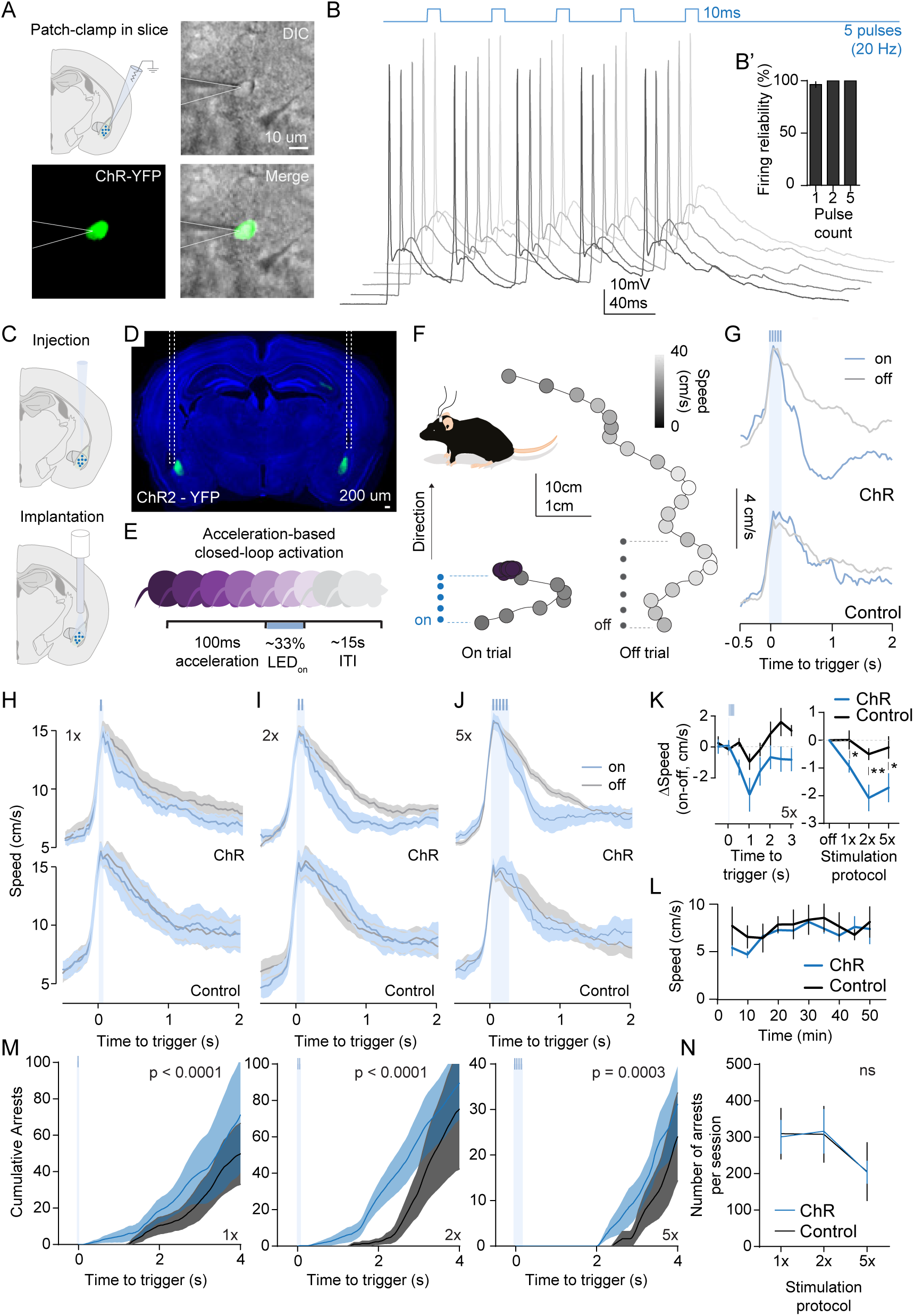
Transient ^NL189^BLA neuron activation decreases movement and promotes arrest. A, Top left, schematic of patch-clamp in slice to test ^NL189^BLA neurons expressing ChR and YFP. Top right and bottom, DIC, fluorescence and merged image showing the patching of a single ^NL189^BLA neuron in slice. B, Top, 5x stimulation protocol. Bottom, effect of five 10ms pulses at 20Hz on neuronal firing (five trials) of a representative ^NL189^BLA neuron. B’, Inset, showing the reliability to elicit firing with one, two and five pulses at 20Hz. Scale: 10mV, 40ms. C, Schematic of a conditional AAV virus expressing ChR2-YFP injection followed by optical fiber implantation. D, Representative coronal section of a mouse bilaterally injected in BLA with the conditional virus expressing ChR2 - YFP and implanted with 200μm optical fibers. Dashed lines underlie the fiber track. E, Acceleration-based closed-loop optogenetic activation protocol. The different colors denote that the animal has to be moving to trigger the trials. F, Top left, scheme of a mouse with bilateral optical fiber implantation attached to patch-cords. Right, two speed color-coded trajectories in light *on* (left) and *off* (right) closed-loop trials (5x protocol). Aligned dots (blue for the on trials and gray for the off trials) showing the triggering of the LED for 5x protocol. G, Speed in closed-loop trials of two representative animals during light on (on, light blue) or off (off, gray) for the ChR (top) and Control (bottom) group. Scale: 4cm/s. I, J, and K, Speed in closed-loop trials using three stimulation protocols (1x, 2x and 5x; n = 7 ChR and n = 5 Control group). K, Left, δSpeed amplitude (on – off trial speed) elicited by the 5x stimulation protocol. Binning: 500ms. Right, δSpeed amplitude (on–off trial speed during 2s from starting the stimulation) elicited by the three stimulation protocols. The ChR group is in blue while the Control is in black. \**p* < 0.05 and **p < 0.01 by unpaired *t-*test for three different stimulation protocols (ChR versus Control group). L, Binned raw speed in ChR and Control animals during the 5x stimulation protocol. M, Left to right, Cumulative arrests after 1x, 2x and 5x stimulation. *p values* are calculated with unpaired two-tailed t-test from 0 to 4s. N, Overall number of arrests in the entire session for the three stimulation protocols. Data are presented as mean ± SEM.

### Inhibition of ^NL189^BLA neuron facilitates movement speed

As the next step, we adopted a loss-of-function optogenetic method using the inhibitory opsin Jaws (Chuong et al., 2014) to silence ^NL189^BLA neurons (Figure 3). First, we tested the photocurrent elicited by light-activation of Jaws compared to eNpHR3.0 using whole-cell somatic patch-clamp recordings in slice (red-light, λ = 630nm; power^max^ = 3.3mW; Figure 3A, B and Figure S9). As we found that Jaws was much stronger in eliciting light-inducing hyperpolarizing photocurrent compared to eNpHR3.0 (Figure S9), we used this inhibitory opsin to further screen its effect on neuronal excitability (Figure 3A-C and Figure S9). The firing of ^NL189^BLA neurons elicited by a step-injection of somatic depolarizing current was completely inhibited by a 2s red-light pulse and immediately restored after the light was switched off (Figure 3A, B). Additionally, inhibition caused a shift of the input-output function of ^NL189^BLA neurons towards more hyperpolarized values (Figure S9). Jaws-induced ^NL189^BLA neurons inhibition rapidly and reliably blocked single action potentials elicited by 10ms-current step (Figure S9). Furthermore, by co-injecting two conditional AAV viruses expressing Jaws and GCamp6f (1:1), we demonstrated the transient silencing of ^NL189^BLA neurons by combining optogenetic inhibition with *in vivo* calcium imaging of ^NL189^BLA neurons using the newly developed *nVoke* miniaturized microscope (Owen, Berke and Kreitzer, 2018; Figure 3C). Conversely to the Chr2 experiments, speed- and acceleration-locked closed-loop optogenetic silencing of ^NL189^BLA neurons (2s red-light) revealed an increase in movement speed in comparison to off trials and Control animals (Figure 3D-G). This set of experiments further confirms that the normal activity of ^NL189^BLA neurons mediates moment-to-moment changes in movement during exploration.

**Figure 3.**
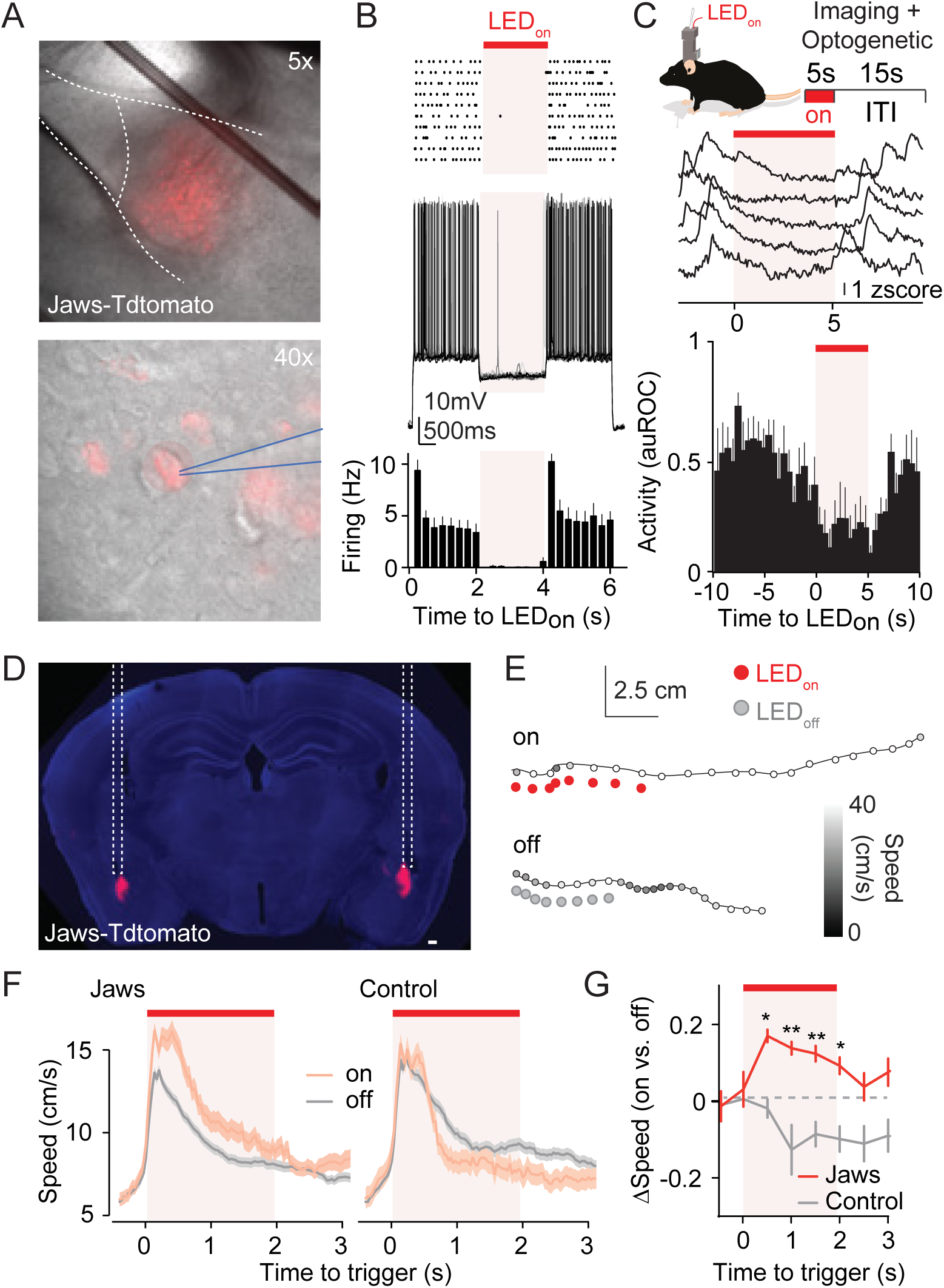
Inhibition of ^NL189^BLA neuron facilitates movements. A, Top, 5x-magnification of a slice in the patch-clamp chamber held by a platinum anchor (black bar) showing the expression of Jaws and Tdtomato in BA. Bottom, 40x-magnification showing the recording from a single ^NL189^BLA neuron in slice. Blue lines show the patch-pipette. B, Top, raster plot of firing elicited by a step of current in a ^NL189^BLA neuron with 2s 630nm-light. Center, firing trace for the raster plot. Bottom, 500ms-binned time-course of light-induced inhibition for all averaged data (ChR, n = 6; Control, n = 5). C, Top inset, schematic of a mouse mounted with an nVoke mini-microscope to simultaneously image neuronal activity and elicit red-shifted opsin activation with 5s-light and 15s of inter-trial interval (ITI). Center, representative neuronal calcium activity trials before, during and after light inhibition *in vivo*. Scale: 1 z-score. Bottom, auROC activity time-course as shown in panel B but for *in vivo* imaging. D, Representative coronal slice using a conditional AAV virus expressing Jaws and Tdtomato. E, Two speed color-coded trajectories in *on* (top) and *off* (bottom) closed-loop trials. The LED_on_ is shown in red circles. As shown in figure 2, triggering of the light is caused by a moment of 100ms-acceleration (when animals are moving), with a variable ITI of ∼15s. F, Closed-loop as for figure 2 but triggering 630nm-light for Jaws and Control (n = 5 for Jaws and n = 4 for Control). G, Change in speed between on (pink) and off trials (gray) for Jaws and Control group in triggering trials. \**p* < 0.05, \*\**p* < 0.01 by unpaired *t-*test within time series of Jaws (red) and Control (black). Data are presented as mean ± SEM.

### Arrest ^NL189^BLA neurons increase in an experience-dependent manner

After assessing the direct involvement of ^NL189^BLA neurons on movement speed and momentary behavioral arrests, we asked whether their activity changes as movement dynamics in exploration change with experience. As previously mentioned, in our emergence exploratory assay, animals freely explore a novel arena starting from a familiar home-shelter on 5 sessions, one per day, from day 1 to 5 (Figure 1A, B). In line with a process of familiarization, exploratory arrests increased across days in the 100cm-arena, while their number is kept constant in the more familiar 20cm-arena (Figure 4A-C). The increase in exploratory arrests was related to the number of exploratory trips (Figure 4D). Voluntary arrests started to appear at the 26^th^ exploratory trip when 75.81 ± 7.485% area explored (Figure 4D, grey dashed line).

**Figure 4.**
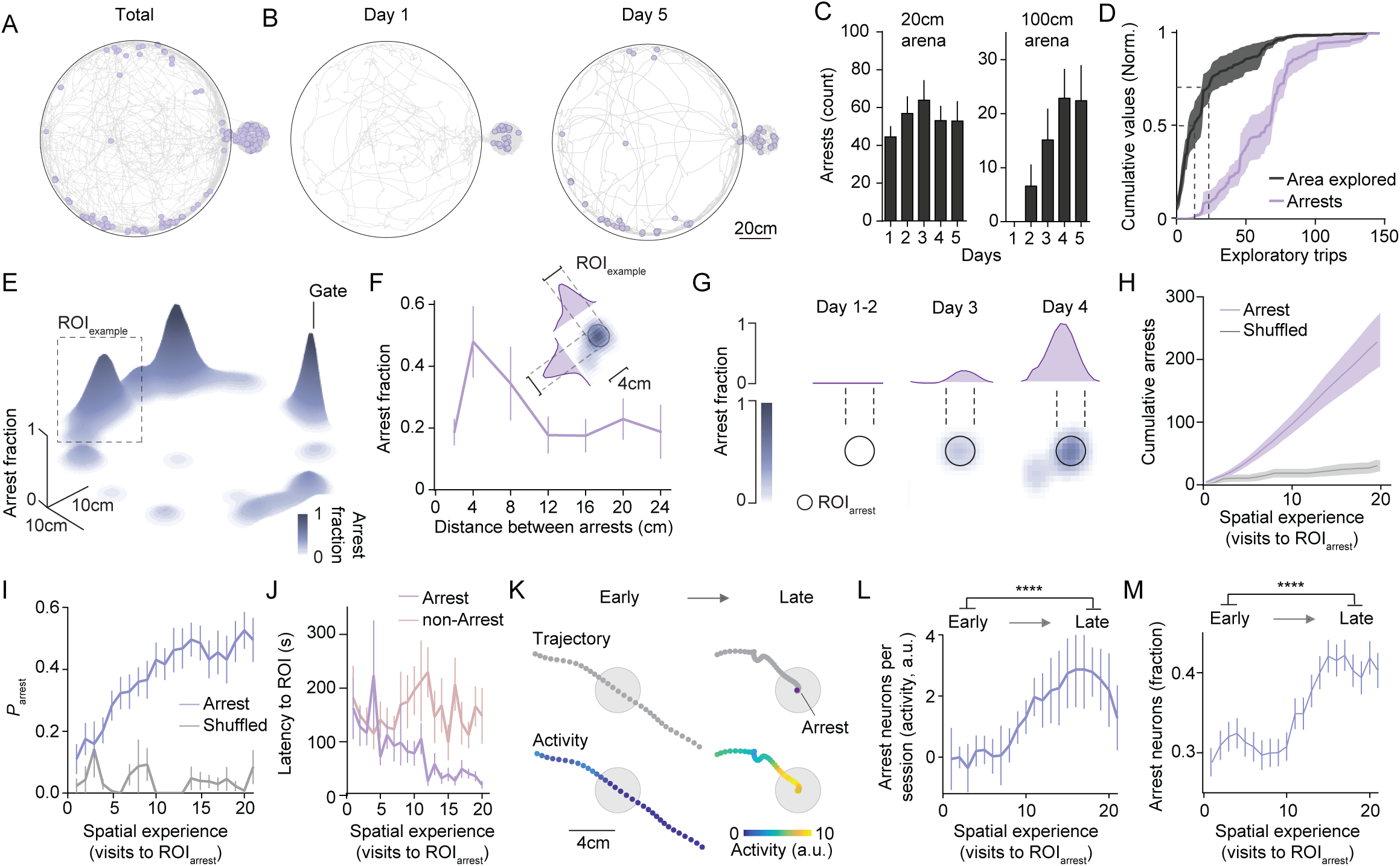
^NL189^BLA neurons encoding self-paced arrest are recruited in an experience-dependent manner in familiar locations. A, Representative aerial view of a mouse’s trajectory super-imposed to arrests (purple circles) for all days. B, Same as A but for days 1 and 5. C, Arrest count in the small (left) and big (right) arena for each day. D, Cumulative area explored (black) and arrests (purple) in each trip (n = 7). At 50% of cumulative area explored there was no detection of behavioral arrests (gray dashed line at 0.5 of cumulative values, normalized). The first occurrence of arrests is denoted with a grey dashed line. E, 3D plot showing the arrest fraction of a representative animal for the five days. An ROI_example_ delineated by dashed line was used in panel F and G. F. Proportion of arrests at different distances. Inset, ROI_example_ showing the distribution of arrests (proportion) for different distances. G, ROI_example_ showing the increase in proportion of arrests across days (from days 1 to 4). H, Time-course of the cumulative arrest count in ROI_arrest_ in relation to the number of visits to that ROI. Shuffled data are shown in the gray time course. I, Probability of arrest (P_arrest_) at an ROI where animals arrest in relation to number of visits to that ROI. J, Inter-entrance interval (IEI, s) to either ROI_arrest_ (areas where mice arrested) or ROI_Non-Arrest_ (areas where mice did not arrest) in relation to number of visits to those places. K, Top, representative animal trajectory in an early and late visit. Purple dot represents the arrest. Bottom, heatmap of a single neuron recorded from the same animal. L, Average δActivity of arrest neurons in relation to number of visits to arrest areas. Early (1-5) and late (16-20) spatial experience are compared. M, Fraction of arrest ^NL189^BLA neurons to be active increases with spatial experience. \*\*\*\**p* < 0.001 by paired *t-*test, early versus late. Data are presented as mean ± SEM.

These data indicate that a certain knowledge of the environment is required before voluntary exploratory arrests start occuring. We therefore investigated if the onset of experience-dependent exploratory arrests was related to the change in representation of the entire arena, or occurred in a spatially-segregated manner, based on the number of visits to specific places. First, we quantified the distance between the arrests’ coordinates (Figure 4F) and found that they accumulate within a distance of 4cm (Figure 4E, F). Second, to understand whether such arrests appeared based on a spatially-segregated experience, we tracked the visits of the animal (referred here as spatial experience) to defined region-of-interest were the animal had arrested at least once (ROI_arrest_; Figure 4G). The diameter of the ROI_arrest_ was chosen based on the most common distance between arrests, 4cm. Interestingly, the cumulative exploratory arrests increased upon entrance to these ROI_arrest_ (Figure 4H). Accordingly, the probability of arrest (*P*_arrest_) increased upon spatial experience in ROI_arrest_ compared to other ROIs of similar size (shuffled data; Figure 4I). Furthermore, the inter-entrance interval (IEI, s) to ROI_arrest_ decreased with number of visits, but it remained constant in ROIs where the animal had never arrested before (ROI_Non-Arrest_; Figure 4J). Additionally, the entrance preference to ROI_arrest_ was higher compared to ROI_Non-Arrest_ (preference ratio: (ROI_arrest_ - ROI_Non-Arrest_)/(ROI_arrest_ + ROI_Non-Arrest_) = 0.3279 ± 0.04162). These results show that exploratory arrests depend on an interaction between spatial and behavioral experience, as mice tend to momentarily arrest in places where they previously arrested (not only visited). Importantly, the fact that they prefer to visit (and arrest) in places where they previously arrested suggest once more that momentary arrest is not a defensive response associated with negative valence, such as freezing.

Next, we sought to understand whether neuronal activity of ^NL189^BLA neurons encoding self-paced arrests was modulated by spatial experience. We sorted arrest ^NL189^BLA neurons by selecting them based on the positive ROC- and anti-CC-based analysis (Figure S6). We found a substantial increase in the proportion of ^NL189^BLA neurons encoding self-paced arrests across days of exploration (Figure S6). The activity of arrest ^NL189^BLA neurons was positively correlated with the daily arrest count (Figure S6). To understand whether the arrests appeared based on fine-spatial experience, we analyzed the overall activity of arrest ^NL189^BLA neurons during early (1-5) and late (16-20) entrances to ROI_arrest_ (Figure 4K-M and Figure S6). As found for behavioral arrests (Figure 4H, I), the neuronal activity of arrest ^NL189^BLA neurons increased with the number of visits to ROI_arrest_ (ROI where they arrest at least once, Figure 4K, L). The fraction of arrest ^NL189^BLA neurons and their probability to be active also increased with experience (Figure 4M). Overall, these findings reveal that ^NL189^BLA neurons encoding self-paced behavioral arrests are recruited in an experience-dependent manner, and tend to fire in familiar locations where animals previously arrested.

### Amygdala silencing impairs experience-dependent exploratory arrest

In order to investigate the necessity of ^NL189^BLA neurons for animals to arrest in locations that they visited and where they arrested before, we performed location-specific optogenetic silencing using Jaws in the emergence exploratory paradigm. During acquisition (day 1 to 5; figure 5A, B), the entrance to one ROI triggered a 2s-activation of a 630-nm LED (ROI_on_). A control area (ROI_off_) located on the opposite side of the 1m-arena was used as an internal control to monitor the general exploration and calculate the place preference. We observed that animals expressing Jaws in BLA arrested less than the controls after crossing the ROI_on_ (Figure 5C). Importantly, this was not because they avoided the area, as the daily number of entrances to this ROI did not differ between the two experimental groups (Figure 5D). In order to understand whether this effect was spatially-restricted to amygdala silencing, we measured the cumulative arrests based on the spatial experience from crossing the ROI_on_ and ROI_off_ (number of visits to ROIs, Figure 5E and Figure S10). In accordance with the previous results, cumulative arrests increased based on spatial experience (Figure 5E). However, cumulative arrests were drastically decreased at the entrance to the ROI_on_ in the Jaws versus Control group, but remained unaffected in the ROI_off_ (Figure 5E and Figure S10). During a probe test (P) administered at day 6, where no light was turned on in the ROI_on_, we observed that the arrest count was initially higher in the Control compared to the Jaws group, but normalized after 25 visits (Figure S10). No significant difference was observed for the entrance to ROI_off_ between Jaws and Control mice (Figure S10). Furthermore, we calculated the preference ratio between the entrances to ROI_on_ and ROI_off_ and found no significant difference between the Control and Jaws group during the probe day 6 (Figure 5F), again suggesting that the manipulation that affected exploratory arrests based on spatial/behavioral experience did not affect the value of the ROI (Namburi *et al.*, 2015; Beyeler, Namburi, Gordon F Glober, *et al.*, 2016; Kim *et al.*, 2016; Beyeler, C.-J. Chang, *et al.*, 2018). Other exploratory parameters were used to quantify possible avoidance/preference or anxiety-like behaviors (Jain *et al.*, 2012) that the silencing of ^NL189^BLA neurons might have induced. No differences were found between the Jaws and Control group in the preference between ROI_on_ and ROI_off_ or the probability to enter the ROI_on_ (Figure S10). Combined, these results show that ^NL189^BLA neurons are fundamental for the development of experience-dependent arrests in defined familiar places where animals have arrested before. On the other hand, this neuronal population does not affect place preference.

**Figure 5.**
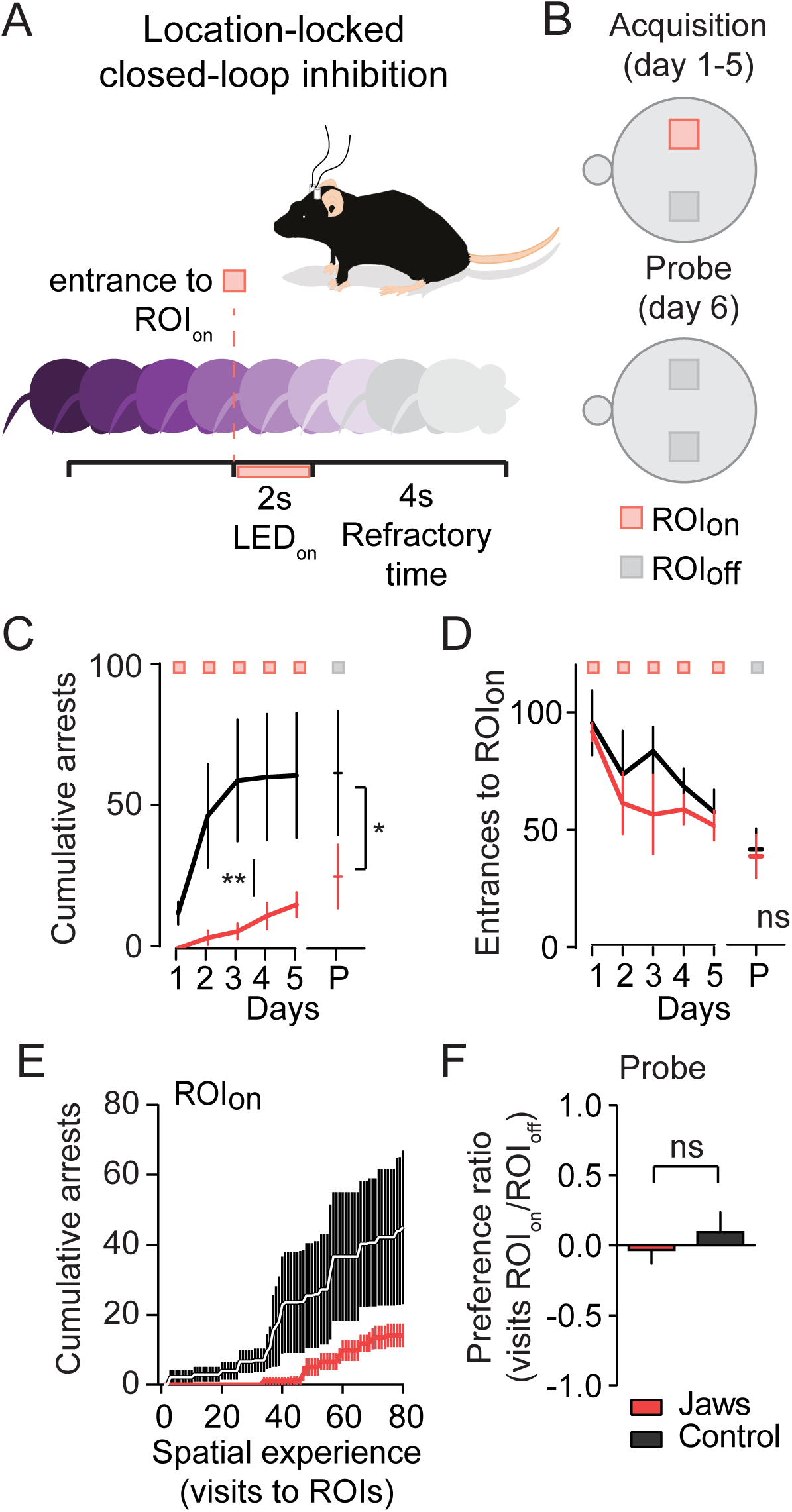
Amygdala silencing impairs the emergence of experience-dependent exploratory arrests. A, Location-based closed loop optogenetic silencing behavioral protocol at the entrance of the ROI_on_. Upon entrance to the ROI_on_, the LED is triggered for 2s 630nm-light delivery followed by 4s refractory time. B, Same as A, Optogenetic protocol executed for the days of acquisition (day 1-5, inhibition in ROIon) and during the probe test (day 6, no inhibition). A control ROI_off_ is used to monitor the avoidance or preference to the ROI_on_. C, Cumulative arrests count from day 1 to 5 for the Control (black; n = 7) and Jaws (red; n = 6) groups followed by a probe trial (P, no light) on day 6. \**p* < 0.05 and \*\**p* < 0.01 by unpaired *t-*test between the two groups for all five days. D, Number of entrances to the ROI_on_ from day 1 to 5 followed by the probe test (P). E, Cumulative arrest number versus number of visits to ROI_on_ during days 1-5. F, Preference ratio between ROI_on_ and ROI_off_ of the number of entrances in day 6. ^ns^*p* > 0.05 by unpaired *t-*test. Data are presented as mean ± SEM.

### CEA-projecting ^NL189^neurons facilitate exploratory arrest

Classical and recent studies demonstrated that electrical and optogenetic stimulation of the main output structure of amygdala, CEA, as well as direct optogenetic stimulation of different neuronal subtypes in this brain area induces unconditional immobility and a decrease in track length (Ciocchi *et al.*, 2010; Li *et al.*, 2013; Botta *et al.*, 2015). Based on this, we investigated if the specific projection of ^NL189^BLA neurons to CEA (CEA-projecting ^NL189^BLA neurons) could directly elicit behavioral arrests. We first confirmed that the fiber density of ^NL189^BLA neurons was higher in CEA than other output structures, such as bed nucleus of stria terminalis (BST) and striatum (Str, Figure S11). Using optogenetic stimulation in slice recordings, we tested the direct connectivity between BLA and CEA neurons located in the medial subdivision (CEm) known to be involved in eliciting immobility, which are also the ventral-lateral periaqueductal gray matter (vlPAG)-projecting neurons (Ciocchi *et al.*, 2010; Tovote *et al.*, 2016; Figure S12). Confirming previous results assessed with electrical stimulation of BLA to CEA (Samson and Pare, 2005), we found that 54.5% of vlPAG-CEm neurons (12 out of 22) receive direct glutamatergic inputs from ^NL189^BLA neurons (Figure S12). As recent studies show that the mesencephalic locomotor region (MLR), fundamental in locomotor initiation, is targeted by afferents from CEA (Roseberry *et al.*, 2016, 2019; Roseberry and Kreitzer, 2017; Caggiano *et al.*, 2018), we examined whether MLR-projecting CEA neurons can receive glutamatergic inputs from ^NL189^BLA neurons using the same optogenetic approach to target vlPAG-projecting in slices. Not only did we find a direct glutamatergic connection from ^NL189^BLA neurons to MLR-projecting CEA neurons located in CEm, but also identified that this connection was significantly stronger than that of vlPAG-projecting CEA neurons (Figure S12). We were able to elicit oEPSCs onto MLR-projecting CEA neurons with a success rate of 100% (n = 22, Figure S12). In addition, the oEPSCs amplitude recorded from this neuronal population was significantly higher than the oEPSCs recorded from vlPAG-projecting neurons (Figure S12). We also tested whether ^NL189^BLA neurons could directly project to vlPAG, specifically to the neurons known to send their afferents to a pontine motor area (Medulla, Mc) important for freezing behavior (Tovote *et al.*, 2016). As no significant connections were found for all 28 neurons tested (Figure S12), these results provide further evidence that there is no strong projection between BLA to vlPAG that could explain the arrests (Tovote *et al.*, 2016; Xu, Krabbe, Schnitzer, *et al.*, 2016). On the other hand, the excitatory efferents from BLA to different CEA neuronal populations were found to be quite robust and be likely candidates to participate in the triggering of arrests (Figure S12; Roseberry *et al.*, 2016, 2019; Roseberry and Kreitzer, 2017; Caggiano *et al.*, 2018).

Using bilateral conditional retrograde viral expression of ChR in CEA and optical fiber implants above BLA, we therefore targeted ^NL189^BLA neurons projecting to CEA during the speed- and acceleration-locked closed-loop optogenetic stimulation (Figure 6A). Similar to the optogenetic stimulation of the entire population of ^NL189^BLA neurons, CEA-projecting ^NL189^BLA neurons induced a transient decrease in speed in the ChR group (Figure 6B-D). This decrease in movement speed was not observed in the Control group (Figure 6C, D) or by stimulating neurons projecting to striatum (Figure S12). Next, we assessed whether location-locked closed-loop optogenetic stimulation of ^NL189^BLA neurons was able to impact the development of experience-dependent exploratory arrests. We found that the number of arrests increased during the acquisition days in ChR versus the Control group (Figure 6E-F). Again, this manipulation did not change the number of entrances to ROI_on_ (Figure 6G). Also, no differences were observed in cumulative arrests, entrances to the ROI_off_ previously ROI_on_ and preference ratio between ChR and Control animals during the probe day test (probe, P) (Figure 6F-H).

**Figure 6.**
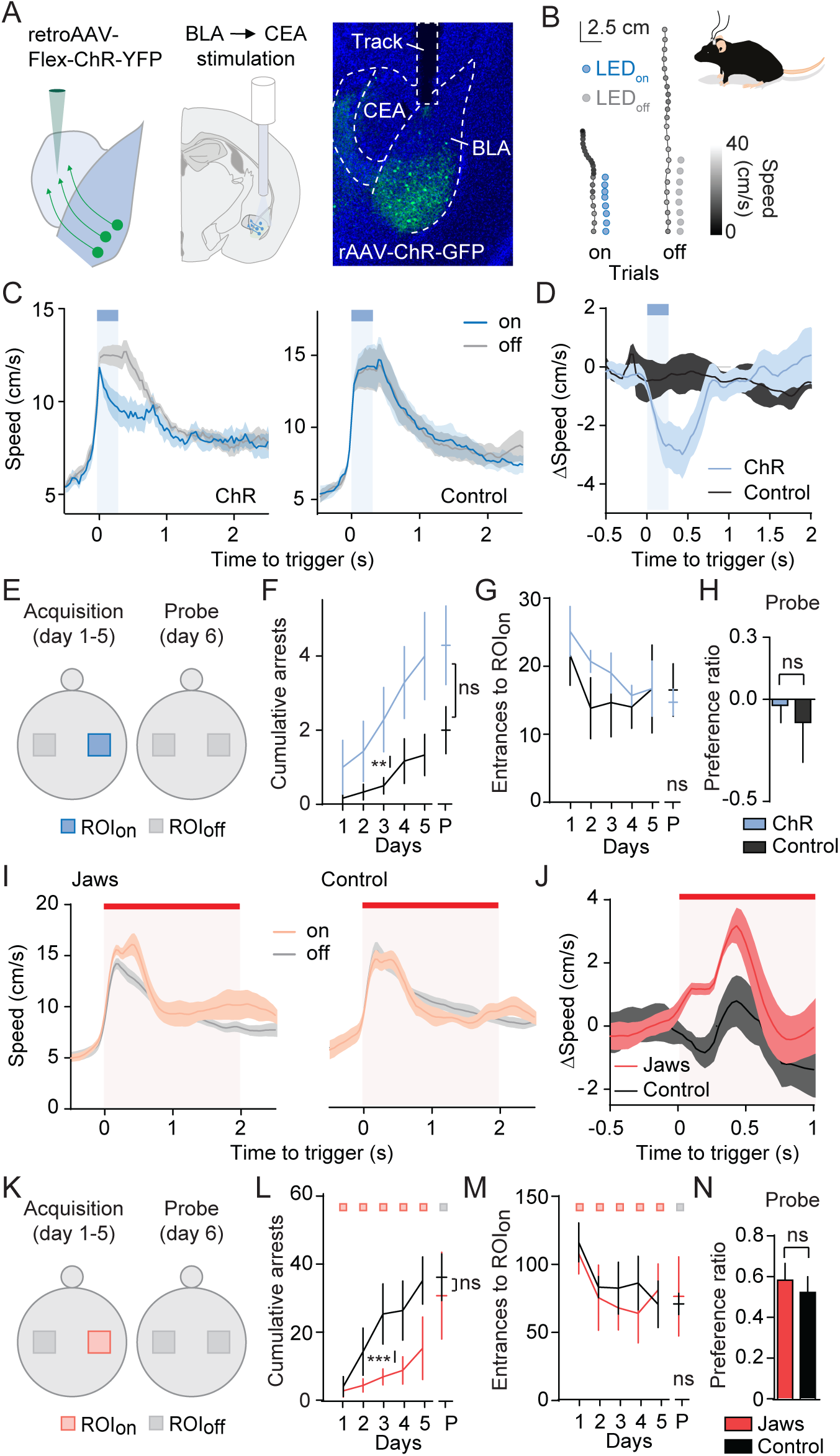
Moment-to-moment arrest is mediated by CEA-projecting ^NL189^neurons. A, Left, injection of a retro conditional AAV virus expressing a floxed version of ChR and YFP. Center, fiber implantation scheme. Right, anatomical location of CEA-projecting ^NL189^BLA neurons and optical fiber implant. B, Two speed color-coded trajectories in *on* (left) and *off* (right) closed-loop trials. Stimulation is denoted by blue circles. Scale bar: 2.5cm, 2.5cm. C, Speed during closed-loop stimulation protocol upon crossing the acceleration threshold for the ChR (left; n = 7) and Control (right; n = 6) group during light on and off trials. D, Change in speed between trials on (light blue) and off (dark) for ChR and Control group in triggering trials. E, Closed-loop location-locked optogenetic stimulation behavioral protocol during acquisition (day 1-5, stimulation in ROI_on_) and probe test (day 6, no stimulation). F, Cumulative arrests count from day 1 to 5 for the Control (black) and ChR (light blue) group followed by probe test (P, days 6). \*\**p* < 0.01 by unpaired *t-*test between the two groups for all five days. G, Number of entrances to the ROI_on_ from day1 to 5 followed by the probe test (P). H, Preference ratio between the entrances to ROI_on_ and ROI_off_ in day 6. ^ns^*p* > 0.05 by unpaired *t-*test. I, Speed during closed-loop stimulation protocol upon crossing the acceleration threshold for the Jaws (left; n = 6) and Control (right; n = 7) groups during light on and off trials. J, Change in speed calculated as the difference between the speed in trials on and off for Jaws (red) and Control group (black) in triggered trials. K, Closed-loop location-locked optogenetic inhibition behavioral protocol in acquisition (day 1-5, inhibition in ROI_on_) and during the probe test (day 6, no inhibition). L, Cumulative arrest count from day 1 to 5 for the Control (black) and Jaws (red) groups followed by day 6 (probe, P). \*\*\**p* < 0.001 by unpaired *t-*test between the two groups for all five days. M, Number of entrances to the ROI_on_ from day 1 to 5 followed by the probe test (P). N, Preference ratio between the entrances to ROI_on_ and ROI_off_ in day 6. ^ns^*p* > 0.05 by unpaired *t-*test. Data are presented as mean ± SEM.

In order to further understand the necessity of CEA-projecting BLA neurons in movement speed and the development of experience-dependent exploratory arrests, we bilaterally injected in CEA a conditional retrograde AAV virus expressing Jaws-GFP (Jaws group) or GFP in Control animals and implanted optical fibers above BLA. Speed- and acceleration-locked closed-loop optogenetic silencing of ^NL189^BLA neurons facilitated movement (Jaws versus Control group; Figure 6I, J and Figure S13), similarly to the manipulation of BLA neurons reported before (Figure 3). Additionally, location-locked closed-loop inhibition of CEA-projecting ^NL189^BLA neurons significantly decreased the cumulative arrests during acquisition, without impairing the number of visits to the ROI_on_ (Figure 6K-M) and the preference ratio between ROI_on_ and ROI_off_ on the probe day (Figure 6N). Our findings demonstrate that CEA-projecting ^NL189^BLA neurons are essential for triggering self-paced behavioral arrests independently of long-term effects on locomotion or place preference.

## Discussion

The results presented here show that a genetically and projection-restricted amygdala neuronal ensemble mediates momentary exploratory arrests that emerge in an experience-dependent manner when animals familiarize themselves with a novel environment. A combination of behavioral analyses, calcium imaging and state-locked closed-loop optogenetic manipulations revealed that ^NL189^BLA neurons control movement speed and behavioral arrest via their projection to CEA. The recruitment of such neuronal population gradually increased in an experience-dependent manner as exploration progressed, primarily in specific areas that animals had visited multiple times, and where they had arrested before. Transient loss-of-function of ^NL189^BLA neurons led to the suppression of momentary arrests, highlighting the fundamental role of this population in experience-dependent arrest. Importantly, these arrests were *not* associated with valence or anxiety-related behaviors. These results unveil a dual role for the amygdala as a novelty/familiarity detector *and* as an effector circuit with the ability to drive or suppress spontaneous movements based on spatial experience during exploratory behavior.

Exploration is required to gain knowledge about a new environment. Although the process of searching has been hypothesized to sometimes be stochastic (Hills *et al.*, 2015), it has been recently identified that an innate strategy, referred to as novelty management, could be responsible for the gradual and well-structured exploration observed in mice (Fonio, Benjamini and Golani, 2009; Thompson, Berkowitz and Clark, 2018). Unrestrained spatial exploratory behavior can be structured into defined trips towards the unknown (Fonio, Benjamini and Golani, 2009). While it had been previously shown that the trajectory of each trip becomes more complex with the progression of exploration, it remained unclear before this study whether specific moment-to-moment actions were shaped by general knowledge of the environment or by recognition of specific spatially-segregated compartments chosen by the animal to initiate further exploration. Our behavioral findings provide evidence that animals choose to momentarily arrest in specific areas that they became familiar with, and where they previously arrested after extensive exploration. These findings are consistent with the novelty management model. Momentary arrests in familiar places can potentially be decision points underlying a “*vicarious trial and error behavior*” (Redish, 2016; Thompson, Berkowitz and Clark, 2018), given that animals initiate and terminate exploratory trips from these locations. In agreement with this hypothesis, it has been shown that a variety of hippocampal neurons encode past, current space and upcoming spatial trajectory sequences during periods of immobility in open field (Kay *et al.*, 2016; Redish, 2016).

One of the challenges with the neural implementation of such a model was to identify neuronal circuits that could simultaneously participate in the processing of novelty/familiarity inputs and also shape movement dynamics. Although it has been hypothesized that the hippocampal formation and other brain regions comprising the medial temporal lobe are fundamental in familiarity recognition (Yonelinas, 2002; Eichenbaum, Yonelinas and Ranganath, 2007; Squire, Wixted and Clark, 2007; Eichenbaum *et al.*, 2010; Sauvage, 2010), they do not seem to be involved in the direct modulation of movements. On the other hand, other independent studies have shown that the amygdala has a direct involvement in contextual learning, novelty and familiarity-detection, but also that it is tightly connected to brain areas involved in defensive freezing responses (Yim and Mogenson, 1989; Wilson and Rolls, 1993; Schwartz *et al.*, 2003; Ciocchi *et al.*, 2010; Farovik *et al.*, 2011; Freeze *et al.*, 2013; Roseberry *et al.*, 2016; Tovote *et al.*, 2016; Roseberry and Kreitzer, 2017). Our study provides extensive evidence that a BLA neuronal population can operate as a neuronal detector-effector circuit, able to integrate experience-dependent contextual information and mediate behavioral arrests.

Inputs into BLA carrying spatial or spatial familiarity signals could potentially originate from the medial temporal lobe, such as hippocampus and perirhinal cortex (Fortin, Wright and Eichenbaum, 2004; Eichenbaum, Yonelinas and Ranganath, 2007; Squire, Wixted and Clark, 2007; Farovik *et al.*, 2011; Tomás Pereira, Agster and Burwell, 2016), which are known to send robust efferents to BLA. The hippocampus and perirhinal cortex could contribute to exploratory behavior, allowing animals to acquire information about novel surroundings based on self-motion (Fortin, Wright and Eichenbaum, 2004; Hines and Whishaw, 2005). BLA neurons may integrate inputs encoding spatial familiarity to guide behavior, and even link spatial experience and the appropriate behavioral output via plasticity, for example via spike-timing long-term potentiation (Jung *et al.*, 2010). The latter could explain why we observed a delayed location-specific arrest formation during the probe day 6 after ^NL189^BLA neurons were inhibited for five days upon entrance to the area (Figure S10). Our data demonstrating that the long-term recruitment of an arrest-encoding BLA circuit originated in familiar locations where animals had arrested before suggest that recognition of spatial familiarity levels can be integrated and processed in the amygdala and used to guide a specific type of exploratory behavior, momentary arrests. These data could also provide a framework to understand the neuronal mechanisms underlying the process of spatial latent learning, through which spatial learning occurs with no apparent reinforcers (e.g. food pellet) or associations (e.g. acoustic tone) (Tolman, 1948).

The implication of the amygdala in mediating experience-dependent behavioral arrests may seem improbable given the attention that it has received in relation to valence and anxiety/fear related behaviors. However, previous independent studies demonstrate that a dose-dependent pharmacological administration of N-methyl-d-aspartic acid (NMDA; NMDA receptor agonist) to BLA triggers stopping behavior, whilst its lesioning decreases immobility (Jellestad and Bakke, 1985; Yim and Mogenson, 1989). Additionally, the amygdala has been revealed to participate in the execution of ongoing actions in humans (Sagaspe, Schwartz and Vuilleumier, 2011). The efferents of ^NL189^BLA neurons to CEA are responsible for the decrease in movement speed and triggering behavioral arrests (Figure 6 and Figure S13). Our results and other groups have found that BLA principal neurons form direct excitatory projections onto a variety of CEA neurons (Samson and Pare, 2005; Ciocchi *et al.*, 2010; Li *et al.*, 2013; Tovote *et al.*, 2016; Figure S12). Fine-tuning of the direct BLA glutamatergic inputs onto defined CEA neurons could elicit arresting behavior via a feed-forward inhibition to midbrain structures known to modulate movement, such as ventro-lateral periacqueductal gray matter (vlPAG, Tovote *et al.*, 2016) as well as the MLR (Roseberry *et al.*, 2016, 2019; Caggiano *et al.*, 2018). Our connectivity studies reveal that ^NL189^BLA neurons have a strong bias to contact MLR-projecting CEA neurons compared to those projecting to vlPAG (Figure S12). This raises the interesting hypothesis of a differential role of parallel circuitries for distinct types of behavioral arrest execution targeted by different contextual inputs. While the vlPAG would be recruited by CEA inputs involved in the generation of strong defensive and learned freezing responses, the hard-wired connection between BLA and MLR-projecting CEA neurons could control the initiation of behavioral arrests unrelated to fear. On the other hand, we cannot exclude that the behavioral arrests elicited by BLA to CEA pathway may be mediated by common long-range projections to locomotor brain structures also recruited during freezing motor responses. In this latter scenario, a shared circuit could be engaged for both freezing and voluntary arrest based on different neuronal computational pattern rules (Botta *et al.*, 2015). Adding to the neuronal circuit complexity, the dorsal striatum may be involved as this brain structure is known to be the main contributor of voluntary initiation, execution and selection of actions (Tecuapetla *et al.*, 2016; Klaus *et al.*, 2017). However, closed-loop optogenetic activation of BLA axonal projections in dorsal striatum did not decrease speed or elicit momentary arrests (Figure S12). Even though we did not find any effect on movement speed stimulating striatum-projecting BLA neurons, it is still possible that amygdala and striatum modulate or compete with each other in order to elicit behavioral arrests at the brainstem level or via CEA.

Although it has been found that stimulation of CEA-projecting BLA neurons induces appetitive and defensive behaviors (Beyeler, Namburi, Gordon F. Glober, *et al.*, 2016; Kim *et al.*, 2016), we did *not* observe that manipulations of ^NL189^BLA neuronal ensemble caused avoidance or preference behavior. This apparent discrepancy may be caused by the genetic background of ^NL189^BLA neuronal population that differs from previous studies. Alternatively or in combination, our specific state-locked optogenetic manipulation differs from the common optogenetic stimulation previously adopted (Beyeler, Namburi, Gordon F. Glober, *et al.*, 2016; Behaviors *et al.*, 2017). This highlights the importance of not simply relying on observational analysis of the results of optogenetic manipulation set *a priori*, but to perform guided state-dependent manipulations to probe the function of ongoing neuronal activity in behavior.

In summary, the data presented here suggests that the BLA-CEA axis acts as a novelty/familiarity detector-effector circuitry for generating self-paced behavioral arrests based on the familiarity of a spatial location. They also reveal circuits mediating a latent learning strategy fundamental to performing efficient and safe exploration of novel surroundings, by which animals move through space to acquire novel information at the interface between the familiar and the unknown.

## Acknowledgments

We acknowledge M. Angelhuber and J. Bauto for the initial consultation regarding behavioral analysis. We thank Helio F.M. Rodrigues for building the behavioral box, Ana Vaz and Mariana L. Correia for mouse maintenance and the Costa lab for helpful feedback. This research project was supported, in part, by the Zuckerman Institute’s Cellular Imaging platform. This project was supported by the Swiss National Science foundation Early Postdoc Mobility, EMBO long-term Postdoc fellowship and NIH K99 NS107721 to P.B. and by the European Research Council Consolidator grant (COG 617142) and the U19 Brain Initiative grant (5U19NS104649) to R.M.C.

## Author Contribution

P.B. designed, conducted the experiments, performed the analysis, discussed the results, and wrote the paper; A.F. conducted optogenetic experiments and histological analysis; A.M.V. and A.M. conducted optogenetic experiments; A.M.V contributed to the MATLAB analysis; L.H. developed the pipeline for automated anatomical reconstruction and analysis, and contributed to histological analysis; C.R.G. provided the transgenic cre mouse line. D.P. discussed analysis; R.M.C designed the experiments and analyses, discussed the results, and wrote the paper.

## STAR Methods

### Contact for reagents and resource sharing

Further information and requests for reagents should be directed to and will be fulfilled by the Lead Contact, Rui M. Costa.

### Experimental Model and Subject Details

#### Mice

All experimental protocols were approved by the Columbia University Institutional Animal Care and Use Committee. All experimental animals were 3 to 5 month-old mice individually housed on a 12 hr light/dark cycle with ad libitum access to food and water. BAC Transgenic mice expressing Cre recombinase under the control of Arhgef6 (Tg(Arhgef6-cre)NL189Gsat/Mmucd, 034805-UCD) were used. The line has been backcrossed onto C57Bl6/J mice for at least 8 generations. Sample size is detailed in the Results or figure legends.

### Method Details

#### Viral injection

Surgeries were performed under sterile conditions and isoflurane (1%–5%, plus oxygen at 1-1.5 l/min) anesthesia on a motorized stereotactic frame (David Kopf Instruments, Model 900SD). Throughout each surgery, mouse body temperature was maintained at 37°C using an animal temperature controller (ATC2000, World Precision Instruments) and afterward, each mouse was allowed to recover from the anesthesia in its homecage on a heating pad. Before starting the surgery animals were intraperitoneally injected with Bupivicane AP (0.5 - 1 mg/Kg). The mouse head was shaved, cleaned with 70% alcohol and iodine, an intradermic injection of bupivacaine was administered and the skin on the skull was removed to allow for aligning of the head, drilling the hole for the injection site and performing the implantation. For the calcium imaging, each animal was unilaterally injected with 500 nl of AAV5-CAG-Flex-GCaMP6f-WPRE.SV40 (titer: 7×10^12^ vg/mL; Addgene) into the left hemisphere of basolateral amygdala (AP: 1.2 mm, ML: 3.4 mm, DV: 4.2 mm) using a Nanojet III Injector (Drummond Scientific, USA) at a pulse rate of 2.6 nL per second, 9.9 nL per pulse every 5s. To avoid the 500 nL of volume being delivered only in one location, potentially causing tissue damage, all basolateral amygdala injections were performed moving the tip of the pipette 50μm deeper than the DV coordinate and injecting a volume of about 100nL every 10μm. After each 10um, the pipette was slowly retracted until reaching the DV coordinate. The injection pipette was left in place for 10 min post-injection before it was slowly removed (rate 200μm/s).

For calcium imaging combined with optogenetic inhibition, the virus AAV5-CAG-Flex-GCaMP6f-WPRE.SV40 was simultaneously injected with AAV-CAG-Flex-Jaws-KGC-tdTomato-ER2 (titer: 1.2×10^13^ vg/mL; Addgene; ratio 1:1). For anatomical studies to assess CamKII and cre co-expression, each animal was unilaterally injected with 500nL rAAV2-CamKII-eYFP (titer: 4×10^12^ vg/mL; UNC) and AAV1-CAG-FLEX-tdTomato (titer: 7.8×10^12^ vg/mL; Addgene) as described above (ratio 1:1; 4×10^12^ vg/mL;). For patch-clamp experiments to characterize the input-output curve, basolateral amygdala was bilaterally injected using AAV1-CAG-FLEX-tdTomato. For connectivity patch-clamp studies, BLA was injected with AAV1-EF1a-DIO-hChR2-YFP (titer: 2.3× 10^13^ vg/mL; Addgene) to express ChR. To target vlPAG-, MLR-projecting CEA neurons and Mc-projecting vlPAG neurons we used a retro AAV-CAG-TdTomato (titer: 1.01×10^13^ vg/mL; Addgene) injected either vlPAG (AP: lamda, ML: 0.6 mm, DV: 2.35 mm), MLR (AP: 4.6 mm, ML: 1.2 mm, DV: 3.6 mm) or Mc (AP: 6.4 mm, ML: 0.95 mm, DV: 5.6 mm). For anatomical experiments in central amygdala, each animal was unilaterally injected (AP: 1.1 mm, ML: 2.5 mm, DV: 4.2 mm) into the left hemisphere using 100nL of a retro AAV virus AAV1-EF1a-DIO-YFP (titer: 3.5×10^12^ vg/mL; Addgene) at a rate of 2.6nL per second, 4.6nL per pulse every 10s. For optogenetic experiments, bilateral injections into basolateral amygdala were performed using 500nL conditional AAV viruses AAV1-EF1a-DIO-hChR2-YFP (titer: 2.3×10^13^ vg/mL; Addgene; ChR group), AAV1-EF1a-DIO-YFP (titer: 3.5×10^12^ vg/mL; Addgene; Control group), AAV-CAG-Flex-Jaws-KGC-tdTomato-ER2 (titer: 7.8×10^12^ vg/mL; Jaws group), AAV1-CAG-Flex-tdTomato (titer: 5.9×10^13^ vg/mL; Addgene; Control group). Using retrograde AAV viruses for the optogenetic experiments to specifically modulate the activity of central-projecting basolateral amygdala neurons, the bilateral injection in central amygdala of AAV-EF1a-DIO-hChR2(H134R)-mCherry-WPRE-HGHpA (titer: 1×10^13^ vg/mL; Addgene; ChR group), AAV-EF1a-DIO-mCherry-WPRE-HGHpA (titer: 6.2×10^12^ vg/mL; Addgene; Control group for ChR), pAAV-CAG-FLEX-Jaws-KGC-GFP-ER2 (titer: 1.3×10^13^ vg/mL; Addgene; Jaws group), AAV pCAG-FLEX-mCherry-WPREv (titer: 6.2×10^12^ vg/mL; Addgene; Jaws-Control group) were performed using the volume of 100nL as stated above. For all the anatomical and patch-clamp experiments, after the injection, the skull was cleaned with saline and the skin sealed with Vetbond tissue adhesive (3M, USA) and stitches (MYCO Medical, USA).

#### Chronic lens and optogenetic implantation

Following the same surgical procedures, about two weeks after viral injection, a 500-nm-diameter gradient index (GRIN) lens (1050-002183, length ∼8.4, NA 0.5, pitch 2; Inscopix, Palo Alto, CA, USA) was implanted in the left mouse amygdala above the injection site (AP: 1.2 mm, ML: 3.4 mm, DV: 4.2 mm). Once in place, the lens was secured to the skull using a combination of black Ortho-Jet powder and liquid acrylic resin site (Lang Dental, USA) as well as super glue (Loctite, LOC1255800). Care was taken to minimize scratches, moreover, the lend was covered with paper/tape to protect its surface. Three weeks after the GRIN lens implantation, the microendoscope baseplate (Inscopix) was mounted onto the head of the mouse under visual guidance using the attached microscope to determine the best field of view. The imaging field of view was inspected and allowed to clear for at least five days prior to imaging and behavioral experiments. For optical stimuli delivery, fiber optics (200 µm diameter, NA=0.66) were implanted 200 um above the site of injection using super glue (Loctite, LOC1255800) and cement (as described above).

#### Calcium imaging in freely-moving animals

Mice were briefly anaesthetized using a mixture of isoflurane and oxygen and the mini-epifluorescence microscope was attached to the baseplate. A period of 20–30 min was allowed to recover in the home cage before experiments started. Fluorescence images were acquired at 20 Hz and the LED power was set at 10–20% (0.1–0.2 mW) with a gain of 4 [excitation: blue light-emitting diode (LED); excitation filter: 475/10 nm, 0.24-0.6 mW/mm2; emission filter: 535/50 nm; Inscopix, Palo Alto, CA]. Image acquisition parameters were set to the same values between sessions to be able to compare the activity recorded. Seven GCaMP6f-expressing NL189-Cre mice were imaged during the emergence exploratory paradigm. The procedure and optical parameters were valid also for the nVoke mini-endoscope (Inscopix) with the only exception that the animal was head-fixed to perform the optogenetic inhibition in basolateral amygdala with Jaws delivering 120 pulses every 10s of 630nm light.

#### Optogenetic manipulation

The power from the optical fibers was 6-7 mW for the 465nm blue light for ChR and 3-4mW for the 630nm red light for Jaws. Before starting the series of behavioral experiments, optogenetic manipulation was tested using slice electrophysiology for ChR and Jaws as well *in vivo* for the case of Jaws. Regarding the Jaws experiments during exploration, we used 6 Jaws animals and 7 Control (3 out of 7 Control animals did not expressed the fluorophore). Regarding the experiments using the conditional AAV virus expressing ChR2 in BLA, all controls expressed the fluorophore YFP. Regarding the six Control animals in retro-ChR experiments, four expressed the fluorophore mCherry in BLA. All Control animals of retro-Jaws experiments expressed the fluorophore mCherry in BLA. All the experimental groups bilaterally expressed either Jaws or ChR.

#### Histology

After completion of the behavioral experiments, mice were transcardially perfused with saline and 4% paraformaldehyde in PBS. Brains were removed for histological analysis and coronal slices were sectioned at 50 μm (Leica vibratome VT1000). For calcium imaging. immunohistochemistry was performed for GCaMP6 expression by incubating the sections with a GFP antibody (GFP Tag polyclonal antibody, Alexa Fluor 488 conjugate, Molecular Probes #A-21311) diluted at 1:1000 in 0.4% Triton-PBS overnight at room temperature. DAPI was used as counterstaining for all the experiments.

#### Imaging

Automated high-throughput imaging of tissue sections was performed on a custom built automated slide scanner using a AZ100 microscope equipped with a 4x 0.4NA Plan Apo objective (Nikon Instruments Inc) and P200 slide loader (Prior Scientific), controlled by NIS-Elements using custom acquisition scripts (Nikon Instruments Inc.). This approach provided images with a lateral resolution of 1mm.

#### Image processing and analysis

Images of tissue sections were processed and reconstructed in ImageJ/Fiji (Schindelin et al 2012) using BrainJ, a collection of custom tools developed to facilitate automated whole brain analysis of tissue sections in a manner similar to serial two photon tomography (Kim et al 2017) and light sheet microscopy (Renier et al 2016). Multi-channel tissue section images are first arranged in an anterior to posterior order and processed to remove external fluorescence and neighboring objects and/or tissue sections. Subsequently, sections are centered and oriented to facilitate a 2D rigid body registration (Thevenaz et al 1998), which ultimately yields a 3D brain volume. In order to detect cell bodies and projections (axons/dendrites), a machine learning based pixel classification approach was employed using Ilastik (Sommer et al 2011) on images that had background subtracted via a rolling ball filter. Several images for each brain were pooled and used for training on cell bodies, projections, and background pixels. The resulting probability images were further processed in ImageJ for segmentation and analysis of cells and projections.

To analyze the resulting data within the context of the Allen Brain Atlas Common Coordinate Framework (CCF), 3D image registration was performed as previously described (Ragane et al, 2012) using Elastix (Klein et al 2010). In this case, DAPI labelling of cell nuclei was used to register the brains to the reference brain using a 3D affine transformation with 4 resolution levels, followed by a 3D B-spline transformation with 3 resolution levels, using Mattes Mutual information to calculate similarity. Following registration, coordinates of detected cells, along with raw image data and segmented projection datasets were transformed into the CCF, allowing analysis of brain region based cell densities, measurements of intensity, and projection densities. Cell plots, density heatmaps, overlays of projections, brain region specific extraction, and subsequent outputs and visualizations were generated using BrainJ and Imaris (Oxford Instruments).

#### Slice electrophysiology

Standard procedures were used to prepare 300-μM-thick coronal slices from 12- to 20-week-old *NL189*-*cre* and *NL189 x Ai9* mice. Briefly, the brain was dissected in high-magnesium (10 mM) ice-cold artificial cerebrospinal fluid (ACSF), mounted on a plate and sliced with a vibrating-blade microtome (V1200S, Leica, USA) at 4 °C. Slices were maintained for 45 min at 37°C in an interface chamber containing ACSF equilibrated with 95% O2/5% CO2 and containing the following (in mM): 124 NaCl, 2.7 KCl, 2 CaCl2, 1.3 MgSO_4_, 26 NaHCO3, 1.25 NaH2PO4, 18 glucose, 0.79 ascorbate. After incubation, slices were left for at least 30 min at room temperature. Recordings were performed with ACSF in a recording chamber at a temperature of 35 °C at a perfusion rate of 1–2 ml/min. Neurons were visually identified with infrared video microscopy using an upright microscope equipped with a 40× objective (Olympus, Tokyo, Japan). Patch electrodes (3–5 MΩ) were pulled from borosilicate glass tubing (G150F-3, Warner Instrument). For current clamp experiments, patch electrodes were filled with a solution containing the following (in mM): 123 potassium gluconate, 12 KCl, 10 HEPES, 10 phosphocreatine, 4 Mg-ATP and 0.3 Na-GTP (pH adjusted to 7.25 with KOH, 295 mOsm). Optogenetic stimulations in slice were performed with a wavelength of 630 nm for Jaws or 465 nm for ChR2 using an optical fiber connected to a PlexBright LD-1 Single Channel LED Driver (Plexon, USA) or using the CoolLED system (pE-300^white^; CoolLED Ltd, UK), respectively. Whole-cell patch-clamp recordings were excluded if the access resistance exceeded 20 MΩ or changed more than 20% during the recordings. Seal resistance, for cell-attached recordings, was around 20 to 50 MΩ and data were excluded if it changed more that 20% from the initial value. Connectivity was assessed by stimulating the axons of BLA for 10ms at 100% power (∼35mW) with 465nm-light for 10ms. Recordings were in presence of TTX (1μM) and 4-aminopyridine (4AP, 100μM). oEPSCs were blocked by further bath-application of the AMPA-receptor antagonist, NBQX (10μM). Successful connected neurons were defined to have oEPSCs >= 10pA with a success rate larger than 50% for all the stimulation trials. Data were recorded with a MultiClamp 700B (Molecular Devices) amplifier and digitized at 10 kHz with a digidata 1550A (Molecular Devices). Lowpass Filter of 0.2 kHz was applied for V-Clamp experiments. Data were acquired and analyzed with Clampex 10.0, Clampfit 10.0 (Molecular Devices) and in-house MATLAB codes. All chemicals for the internal and external solutions were purchased from Fluka/Sigma (Buchs, Switzerland). Glutamatergic blockers were purchased from Tocris Bioscience (Bristol, UK).

#### Behavioral setup

The dimensionality emergence assay consists of two opaque white cylindrical arenas of 20 cm and 100 cm of diameter with walls 30 cm of height. The opacity avoids reflection of light ensuring a better tracking of the animal without background contaminations. The arenas are interconnected with a 6 cm wide opening that can be closed with an automatic gate moving at 90 degrees. The setup is located inside a sound-attenuating box (2 × 2 × 2m; skeleton in aluminum, Item 24; 25mm 75% Sound-Absorbing foam Sheet, McMaster; built in-house) that also confers the ability to control the general light intensity (∼6 Lux) without creating obvious shades inside the open field. The locomotion of the animal was monitored with a camera (Grasshopper3, GS3-U3-41C6M, Flycapture) with wide-field objective (MVL8M1 - 8 mm EFL, f/1.4, for 1 C-Mount Format Cameras, THORLABS) mounted about 150cm above the two arenas. Spatial resolution was 1 cm per 5 pixels and temporal resolution was 30 frames per second. The animal’s center coordinates (centroid) were acquired online with the open source visual language BONSAI (https://bonsai-rx.org/). The triggering of closed-loop optogenetic stimulation or inhibition was also performed with BONSAI.

#### Exploratory behavioral paradigm

Mice were given four weeks to recover from surgery. Afterwards they were handled for two consecutive days (five min each day) using a dummy endoscopic that mimics the weight of the miniscope or with optical patch cables to habituate them to the calcium imaging or optogenetic behavioral procedures, respectively.

##### Calcium imaging

For the calcium imaging experiments, the emergence assay exploratory procedure was performed in five days. In day 1, the animal was placed in the 20cm-arena for 30 minutes to promote familiarization. Afterwards, the automatic gate opened giving the animal the possibility to explore the 100cm-arena for 20 minutes. From day 2 to 5, the gate opened after ten minutes as familiarization to this area has already occurred in day 1.

##### Optogenetic manipulation

In the emergence assay, the location-specific optogenetic manipulation was made by setting one region of interest (ROI; 20 × 20 cm) in the 100cm-arena. The side of the ROI was counterbalanced between animals and selected to trigger the PlexBright optogenetic stimulation system LED module (Plexon, USA) to illuminate 465 nm blue light or 630 nm red light in basolateral amygdala. For the first five days (acquisition), the position of the ROI was maintained constant to ensure the acquisition of spatial memories. In day 6 (probe test), the entrance to the previous ROI_on_ would not trigger the LED. Inhibition occurred for 2 s upon entrance to the ROI with a refractory time of 4s in which the optogenetic LED could not be triggered. Regarding location-locked closed-loop optogenetic stimulation, it occurred using five 10ms pulses at 20 Hz followed by a refractory time of 4s. In the speed- and acceleration-locked closed-loop experiments, optogenetic stimulation (20 Hz, 5 pulses of 10 ms; 250 ms length) or inhibition (2 s constant inhibition) triggered by animals’ acceleration was performed in the 100cm-arena. In the acceleration protocol, to make sure the animal was moving prior optogenetic stimulation, LEDs were triggered with a probability of about 33% only if the animal accelerated for at least 100 ms with a speed higher than 2 pixels per second (0.2 cm/s). The optogenetic manipulations were repeated for 50 trials. There was a refractory time of 20 s in between optogenetic light stimulation or inhibition. Control trials (trial off: ∼66%) were triggered by the acceleration but did not induce any light-emission.

#### Quantification and Statistical Analyses

##### Arrests detection

Segmentation of the smoothed path into progression segments and lingering episodes was done using the EM algorithm with a two-gaussians mixture model as described (Golani, Benjamini and Eilam, 1993; Drai, Benjamini and Golani, 2000). Briefly, we estimated the amount of motion within a temporal window of 500 ms and determine a threshold value under which a data point will be counted as arrest. We therefore computed the standard deviation (SD) of the distances between the data points and their mean using a sliding window of 0.5s. This value was computed assuming the minimum amount of arrest that the animal can do (Drai, Benjamini and Golani, 2000). The arrests are found by studying the statistical distribution of SD and determining a threshold value. To estimate the distribution of the log max SD values of all the episodes in a session, a density estimator was used. Based on the notion that arrests segments are characterized by low SD and estimated background noise, we used the cut-off of three SDs to have a clear-cut between arrests and movement segments.

##### Spatial analysis of arrest areas

We considered arrests occurring only in the 1m-arena to study their formation in novel environments. For the mouse’s *xy* coordinates collected during calcium imaging studies, we combined the position of the animal within five days of exploration. Second, a spatial analysis was used to detect the first arrests occurring in specific region-of-interests (ROIs) of the big arena. Each first arrest was assigned to a defined round ROI (10 pixel radius, 2 cm radius). If arrests occurred within that radius range from the first arrests, they were counted as belonging to the same area. The script detected multiple spatially-segregated arresting areas that we could follow in time with the advantage that entrances, number of arrests and neuronal activity into these areas could be studied. If the entrances occurred less than 3s apart, they were excluded to avoid a misleading overlap of neuronal activity from two consecutive visits. We excluded first arrests occurring less than 2cm from the border of an adjacent arrest area. Novel entrances were considered from the visit 1 to 5 while familiar entrances from 16 to 20.

##### Preference analysis

Place preference was calculated either using the number of entrances or duration to the ROIon (20 × 20 cm square; LED switched on upon entrance during acquisition) versus the ROIoff (20 × 20 cm square; located in the opposite site from the ROIon; the LED was off upon entrance), using the following formula: *(ROIon-ROIoff) (ROIon+ROIoff)*

##### Computational analysis of neuronal activity

All calcium movies were initially preprocessed in Mosaic (v.1.1.2, Inscopix) for spatial binning (4 × 4 pixels) and motion correction and subsequently analyzed using MATLAB. One-photon imaging is known to contain significant background signals arising from out-of-focal plane light and neuropil (Pnevmatikakis *et al.*, 2016; Klaus *et al.*, 2017; Zong *et al.*, 2017) due to the fluorescence excitation of a relatively large three-dimensional volume compared to, for example, two-photon imaging. A constrained non-negative matrix factorization for endoscopic data (CNMF-E; Pnevmatikakis et al., 2016; Zong et al., 2017) was employed to correct somatic calcium transients. It allowed a superior estimation of calcium transients by eliminating the contaminated background fluorescence as a result of the modeling of spatial and temporal background statistics superimposed to the neuronal signals.

##### Cross-correlation analysis

We calculated the speed of the animal using animal coordinates and convert for time (30Hz) and space resolution (1cm is 5pixels).

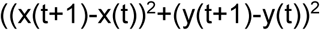

In order to understand the relationship between neuronal activity and speed, we computed the cross-correlation using the following MATLAB function as described (Costa *et al.*, 2006) either using the mean neuronal activity from each animal or single cell:

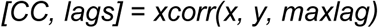

where CC is the cross-correlation of two discrete-time sequences, x and y; lag is a vector with the lags at which the correlations are computed; x is the normalized z-score of the neuronal activity from single neurons or from the entire population; y is the normalized z-score of the animal’s speed; maxlag represents the time-lag range from -maxlag to maxlag (20s). The cross-correlation function was considered statistically significant when more than nine consecutive bins within the [-5 s, 5 s] interval lay outside the [minimum, maximum] bound of its values at intervals [26 s, 40 s] and [0 s, -14 s].

##### Area under the receiver operating characteristic curve (auROC) analysis

For this analysis, we used a method similar to the one described previously (Cohen *et al.*, 2012; da Silva *et al.*, 2018). To produce the ROC curves, we compared the raw calcium activity average during the transition from movement to arrests of each 100-ms bin to the baseline calcium activity (–5 to –1 s before arrest) across trials. The auROC for each bin was calculated using trapezoidal numerical integration. In order to select arrest-responsive neurons, each auROC bin-values were compared to the baseline. Significance was established if at least three consecutive bin-values between the time range 1-2s exceeded 1.5 SD (arrest active neurons) or were less than -1.5 SD (arrest inhibited neurons) from baseline average. After the selection of neurons using two methods, auROC and cross-correlation analysis, arrest active | anti-CC neurons were considered the “pure” arrest neurons and used the spatial analysis of arrest areas. Neuronal activity of arrest active | anti-CC neurons was averaged for each animal and the peak response (difference between the peak between 5 to7s and baseline range, –5 to –1s) was tested during entrance to the novel and familiar areas.

##### Statistical analysis

Mean ± standard error of the mean (SEM) was used to report statistics if not indicated otherwise. For all within-subject quantifications, we calculated the average across all 5 recording sessions. Statistical tests used and the sample size for each analysis is listed in the Results or figure legends. Both parametric and non-parametric tests were used wherever appropriate. Hypothesis testing was done at a significance level of p = 0.05. No statistical methods were used to predetermine sample size. All analyses were performed in MATLAB (MathWorks) while the graphs and statistical tests with GraphPad Prism 6 (GraphPad software, USA). Experimental data were excluded based on imaging quality due to movement artifact, a lack of cells, a bad focal plane, lack of viral injection in the targeted areas and misplacement of the optical fibers.

## Supplementary videos

Movie S1. Example of a mouse performing multiple exploratory trips.

Movie S2. Representative 3D trajectory representing the XY coordinates of the mouse’s centroid versus time (Z-axis; unit: number of frames) before and after gate opening.

Movie S3. Viral expression of tdTomato in BLA. This movie shows the density of viraly targeted NL189BLA neurons expressing tdTomato (color coded heatmap), within the brain (grey, Allen Brain Atlas template brain).

Movie S4. Location of NL189BLA neuron soma (dots) in different parts of BLA (color-coded), within the brain (grey, Allen Brain Atlas template brain).

Movie S5. Calcium imaging of ^NL189^BLA neurons in freely-moving animals (raw signal, left; denoised signal, right).

Movie S6. Example of one momentary arrest occurring at about 3.5s from the start of this movie. Arrest are followed by head scanning of the open field.

Movie S7. Example of other two momentary arrest occurring at 1s (first arrest) and between 2-3s (second arrest) from the start of this movie. The arrests are followed by head-scanning and a change in trajectory’s direction.

Movie S8. Optogenetic activation of ^NL189^BLA neurons during movement. Top left, LED_on_ is triggered at time X of this movie. Activation of ^NL189^BLA neurons causes a momentary arrest.

## Supplementary Information

**Figure SI 1. Behavioral analysis of exploratory trips and locomotion segmentation.** A, Left, representative animal trajectory (*xy axis*) plotted against time (*z axis*) and color-coded speed (heatmap) five minutes before gate opening in the 20cm arena. Right, Same as left but first five min after gate opening. Gate location is shown by the dashed black line and red-filled dot. B, Daily trip count (black bars) and normalized values from day 1 (gray line). C, Quartile trips duration in day 1 and day 5. D, Duration preference in the big versus the small arena from day 1 to day 5. E, Representative trips from a single animal in day 1. F, Similarity of the path (overlap of single trajectories with a range of 2.5cm) during single trips averaged for each single day. Trip similarity is calculated from the first trip. G, Maximum Standard deviation count (SD^max^ count) versus the log of SD^max^ for each segment speed. Arrests segments are denoted by the purple line and dashed line with a threshold speed of 1 cm/s. H, Time course of locomotor speed. Purple bars show the arrests. I, The segmentation with two-gaussians mixture model algorithm finds other behavioral states in movement speed. We use only the transitions from state 2-5 (movement) to state 1 (arrest). Data are mean ± SEM.

**Figure SI 2. Anatomical localization of** ^**NL189**^**BLA neurons.** A, Raw, pixel classification and segmentation image of neuronal ^NL189^BLA soma detection in BLA using a machine learning algorithm. B, Low-magnification of an entire brain slice after unilateral injection in BLA of a conditional AAV virus expressing Tdtomato in ^NL189^BLA neurons. C, medial-lateral view of the BLA and ^NL189^BLA soma reconstructed in 3D. D, same as C but showing the antero-posterior view of BLA for the soma location (left) and the heatmap of soma density of ^NL189^BLA neurons. E, Fraction of ^NL189^BLA neuron soma in three dimensions. F, Fraction of ^NL189^BLA neurons in three part of BLA anterior, posterior BA and LA. Lateral and medial side are shown by the gray and black bars. Data are mean ± SEM.

**Figure SI 3.** ^**NL189**^**BLA neurons correspond to principal cells.** A, Left, Schematic representation of viral injection in BLA of two AAV viruses: one expresses Tdtomato in NL189 neurons (red) while the other GFP in CamKII neurons (green). Center, high magnification of BLA showing the expression of Tdtomato in NL189 (red), GFP in CamKII (green) with an overlap between the two neuronal types (yellow). Right, lower magnition of the injected area (n = 3 animals). B, Summary of the fraction of BLA neurons expressing the fluorescent marker. C, Voltage traces recorded from the soma of NL189 expressing a fluorescent marker in CC (current clamp) mode of three distinct neuronal populations (low, regular and burst firing). Scale: 20 pA, 200 ms. Middle and lower trace shows the first action potential elicited by the current step (marked with a gray bar). Scale bar: 10 mV, 5ms. D, Summary of the inter-spike interval (ISI) distribution in the three neuronal subtypes. E, Summary of the fraction of ISI for the three neuronal populations. F, Left, First ISI versus current step injection (20pA current step) for the three electrophysiological subtypes. Right, Same as left side but for the entire firing frequency elicited by the current injections (n = 35 neurons total). G, Pie chart of proportion of regular, burst and low firing neurons in the ^NL189^BLA neuron population. Negative ^NL189^BLA neurons showed two extra-types of neurons (n = 35). Data are mean ± SEM.

**Figure SI 4. One-photon calcium imaging recordings from freely-moving animals.** A, schematic of viral injection and lens implantation. B, Representative unilateral lens implant. C, Schematic showing the GRIN lens location in 7 animals. D, Raw calcium transient traces from few neurons recorded. Scale: 50 a.u., 2s. E, Neurons count where their activity was recorded from day 1 to 5 (n = 7 animals). Data are mean ± SEM.

**Figure SI 5. Characterization of momentary arrests.** A, Centroid speed at the time of arrest and start of movement (n = 7 mice). Arrest is denoted by the violet bar. B, Angular speed of the head versus the body at the time of arrest and start of movement (n = 7 mice). Inset, Angular speed was calculated from the angle between body (Center-Tail) and Head. C, Speed of the animal prior and during the arrest. C’, Bar graph showing the daily latency to arrest from the maximum speed to time to arrest (0s). \**p* < 0.05 and \*\**p* < 0.01 by One-Way ANOVA followed by Bonferroni *post-hoc* test. D, Arrest duration versus spatial experience in the big arena. Data are mean ± SEM.

**Figure SI 6. Detection of arrests neurons using a variety of methods.** A, Fraction of ^NL189^BLA neurons sorted with a CC analysis from day 1 to 5 for the total number of neurons. B, Left, sorting of different neuronal types based on the ROC analysis (n = 1,435) for the big arena. Right, linear regression between the fraction of arrest neurons and the arrest count in the big arena. C, Fraction of anti-CC and ROC-based arrest neurons each day of exploration (in the big arena). D, Representative ^NL189^BLA neuron showing detected calcium transients (top) overlapping with the z-scored speed (center). Bottom, pie chart of the fraction of neurons active during deceleration in day 1 (left) and 5 (right). E, Summary of the z-scored speed when the neurons in day 1 (left) and 5 (right) were active (time to neuronal calcium transient at 0s). F, Sorting of the different neurons active during deceleration (z-scored speed < -0.5 between -1 and 1 from 1s baseline, dark trace) and not significant (z-scored speed > -0.5, gray). G, Bar graph of the δActivity in early (left) or Late visits to ROIs (right) during arrest (arrest, light violet bar) or movements (move, white bar). \*\*\**p* < 0.001 by unpaired t-test. H, Probability of ^NL189^BLA neurons to be active, respectively, increases with spatial experience. J, Before-after graph bar of the fraction of active arrest ^NL189^BLA in early (left) or late visits to ROIs (right) during arrest (arrest, light violet bar graph) or movements (move, white bar graph). \**p* < 0.05 by paired t-test. Data are mean ± SEM.

**Figure SI 7. Channelrhodopsin-activation protocols to mimic physiological firing.** A, Single 10 ms light activation square pulse (465nm, 1x protocol) on a representative NL-189 neuron in BLA. Scale:10 mV, 40 ms. Inset, voltage trace shows the reliability of the activation. Scale: 10 mV, 4 ms. B, Same as a but using 2 pulses stimulation (2x protocol). C, Same as A and B but using 5 pulses (5x protocol). D, From left to right, bar graphs summarizing the firing reliability, optogenetic evoked action potential count (oAPs count), latency to spike, jitter and resting membrane potential (V_rest_). Data are mean ± SEM.

**Figure SI 8. Effect of multiple** ^**NL189**^**BLA neuron optogenetic stimulation protocols on speed and avoidance.** A, B, and C, 1x, 2x and 5x activation protocol on ChR (top) and Control group (bottom) on normalized speed triggered by acceleration in ∼30% of trials (on, blue line) and off (gray line). b and c, same as panel a but for 2x and 1x protocols, respectively. D, Top, difference in speed between on and off trials for the three stimulation protocols in ChR (blue traces) and Control group (black traces). E, Time course of the normalized speed for the entire experimental optogenetic session. Speed was normalized by the first two minutes baseline before the acceleration-based closed loop experiment started (referred as Optogenetic Protocol). F, Distance from center at the time of blue light on for Control (black) and ChR (light blue) group. G, Normalized distance from center during the 5x closed-loop stimulation protocol. Data are mean ± SEM.

**Figure SI 9. Activation of Jaws induces strong and reliable neuronal inhibition.** A, Representative traces showing the effect of Jaws activation with 1s square pulse of 630nm-light at 100%, 50% and 25% power recorded from ^NL-189^BLA neurons in Voltage-Clamp (VC, -70mV; Top) and Current-Clamp (CC; bottom) mode in slice. Scale bar: top, 20pA; bottom, 5mV. 100% power correspond to 3.3mW. B, Left, photocurrent elicited by Jaws (red line) and halorhodopsin (eNpHR3.0, black line) activation at different powers recorded in VC mode at -70mV (n = 10, Jaws; n = 6, eNpHR3.0). Right, photo-hyperpolarization induced by Jaws or eNpHR3.0 activation at three different powers in CC mode (n = 12, Jaws; n = 6, eNpHR3.0). C, Top row, baseline voltage traces elicited by current square pulses (180, 280 and 380 pA) recorded from a single representative ^NL-189^BLA neuron. Scale: 10mV, 200ms. Bottom row, same neuron as panel A but elicing the firing with square current pulses together with Jaws activation at 100% power. The current injection is shown by the square black pulse while optogenetic inhibition is applied by giving a square pulse (red). D, Current injection in CC mode from the same neuron in panel A-C during the current pulse of 1s length. Each dot represents an action potential elicited by the current injection (I_injection_). Baseline is in gray while current injection combined with Jaws activation is shown in red. E, Rheobase, first current step able to elicit one action potential, in baseline versus Jaws activation. ***p<0.001 by paired test (n = 11). F, Input-output function of ^NL-189^BLA neurons in baseline and adding Jaws activation (n = 11). G, Representative traces showing two consecutive 10ms current steps of 650pA able to elicit one action potential in Control (630nm-light off, left trace) but not in Jaws (630nm-light on, right trace) condition. Bottom left, same trace as the top row of panel G showing the protocol to assess the reliability of Jaws activation. H, Reliability of the current injection in order to elicit action potentials in Control as well in Jaws condition. ****p < 0.0001 by paired t-test (n = 10). Data are mean ± SEM.

**Figure SI 10. Silencing of** ^**NL189**^**BLA neurons does not affect many exploratory behavioral parameters.** A-I and K, M, O, Time course of different parameters associated with exploratory behavior and arrest for the Jaws (red; n = 6) and Control group (black; n = 7) from day 1 to 6. Day 6 is the probe day (P). Pink squares denote the days of optogenetic inhibition upon entrance to the ROI_on_ while gray squares show the probe day 6 (ROI_on_ become ROI_off_; P). J, Preference ratio calculated using the duration in ROI_on_ and ROI_off_ during the probe day 6, in the absence of optogenetic silencing. ^ns^*p* > 0.05 by unpaired t-test between Jaws (red) and Control (black) group. L, Cumulative arrests versus spatial experience (defined as entrance to the ROI control, ROI_off_). N, Before and after graph showing the arrests count after crossing ROI_on_ during light on and off condition in Jaws (left, *p =* 0.0002 by two-tailed paired *t-*test) and Control group (right). P, Left, Normalized cumulative arrests versus entrance to the previous ROI_on_ from entrance 1 to 25. \*\*\*\**p* < 0.0001 by unpaired *t-*test from ROI entrance 1 to 20 (Jaws versus Control group). O, Normalized cumulative arrests versus entrance to the previous ROI_off_ in the probe day6. ^ns^*p* > 0.05 by paired *t-*test from ROI entrance 1 to 20 (Jaws versus Control group). Data are mean ± SEM.

**Figure SI 11. Machine learning-based detection of axonal fibers from** ^**NL189**^**BLA neurons.** A-C, Top, raw images with inverted color of the axonal fibers of ^NL189^BLA neurons. The conditional AAV virus expressing Tdtomato was injected into BLA. Bottom, Probability of axnal projection automatically detected by in-house developed machine learning-based algorithm. D, Normalized quantification of the fiber density in BLA as well as in CEA, BST and Str for animal injected with a conditional AAV virus expressing Tdtomato in BLA. E, Same as D but for animals injected in CEA with a conditional retro-AAV expressing YFP to target CEA-projecting BLA neurons. Data are mean ± SEM.

**Figure SI 12. Direct glutamatergic connection between** ^**NL189**^**BLA and brainstem-projecting CEA neurons.** A, Scheme of an unconditional retro-AAV virus injected in vlPAG (left) or magnocellular nucleus (Mc, middle) or the locomotor region (MLR, right) to retrogrately express Tdtomato and target projecting neurons. B, Scheme of the viral injection for the conditional expression of ChR in ^NL189^BLA neurons. C, Injection site showing the virus (panel A) injected in vlPAG. Right, Expression of Tdtomato in vlPAG-projecting CEm neurons. D, Merged DIC and YFP image showing the viral expression of the conditional AAV virus expressing ChR and YFP (as panel B). Patch-pipette is shown by the blue lines. Recordings were performed in vlPAG from Tdtomato-expressing neurons (right side). E, Optogenetic stimulation protocol of the ^NL189^BLA neuron terminals in the medial part of CEA combined with patch-clamp recordings from vlPAG-projecting CEA neuros in slice (left). Middle, Optogenetic recordings from Mc-projecting vlPAG neurons. Right, Same as left but for MLR-projecting CEA neurons. To assess monosynaptic connectivity from BLA to CEA, each patch-clamp recording was performed in presence of tetrodotoxin (1μM) and 4-aminopyridine (100μM) to block voltage-gated Na^+^ channels and K^+^ channels, respectively (Petreanu and Svoboda, 2009). F, Summary of optogenetic stimulation of the afferents of ^NL189^BLA neurons that evoked excitatory postsynaptic currents (oEPSCs) at time 0ms by a 10ms blue light pulse (465nm) in the connected (darker pink trace average, n = 12) versus the non-connected (gray, lack of oEPSCs, n = 10) vlPAG-projecting CEA neurons. Pie chart shows the fraction of connected (dark violet) versus non connected neurons (light gray). Middle, Recording from Mc-projecting vlPAG neurons where no significant connections were found (n = 28). Right, oEPSCs recorded from MLR-projecting CEA neurons elicited by light stimulaton of ^NL189^BLA neurons’ axons (n = 22). The pie chart shows 100% connections (dark blue). We considered successful connections with an amplitude ≥ 10 pA that occur more than 50% of the trials. The holding voltage (V_hold_) was at -80 mV. G, oEPSC amplitude average for all projecting neurons. Each gray circle represents the oEPSC recorded from a single neuron. ##*p* < 0.0035 by unpaired *t-*test; \*\**p* < 0.01 and \*\*\*\**p* < 0.0001 by One-Way ANOVA followed by Tukey’s Multiple Comparison Test. H, oEPSC latency (ms) for vlPAG- and MLR-projecting CEA receiving ^NL189^BLA neuron inputs. \*\**p =* 0.0028 by unpaired t-test. Success rate to elicit successful oEPSCs (amplitude > 10pA) in the connected versus non-connected CEA neurons. I, Before-and-after bar graph showing oEPSCs evoked by 10ms light pulse during baseline and adding 10μM NBQX in the patch-clamp chamber for vlPAG- (left; n = 10) and MLR-projecting CEA neurons (right, n = 15). \*\*\**p* < 0.001 by paired-test. J, Schematic of speed- and acceleration-locked close-loop optogenetic stimulation of dorsal striatum (Str) afferents from BLA. K, Speed profile at time of BLA to Str stimulation for ChR and Control groups (n = 5, ChR; n = 4, Controls). Data are mean ± SEM.

**Figure SI 13. Inhibition of CEA-projecting** ^**NL189**^**BLA neurons the development of momentary arrests.** A, Effect of the acceleration-locked closed-loop optogenetic inhibition of CEA-projecting ^NL189^BLA neurons (on trials, pink) versus off trials (gray) for the Jaws (left) and Control group (right) on normalized speed. B, Change in normalized speed calculated as the difference between the normalized speed in trials on and off for Jaws (red) and Control group (black) in triggering trials. Enlarged (left) and high magnification (right) of the effect of light inhibition in the Jaws group the first 500ms from the triggering of the LED. Data are presented as mean ± SEM.

**Figure SI 14. Histological analysis of viral expression and optical fiber implant location.** A, Sections show the implantation of the optical fibers. B, Heat Map shows the soma density for the experimental groups that were used for the optogenetic experiments (from left to right: conditional AAV virus expressing Jaws and ChR in BLA as well as the conditional retro virus injected in CEA expressing ChR and Jaws in BLA).

