## Supplementary Figures for "An amygdala circuit mediates experience-dependent momentary exploratory arrests"

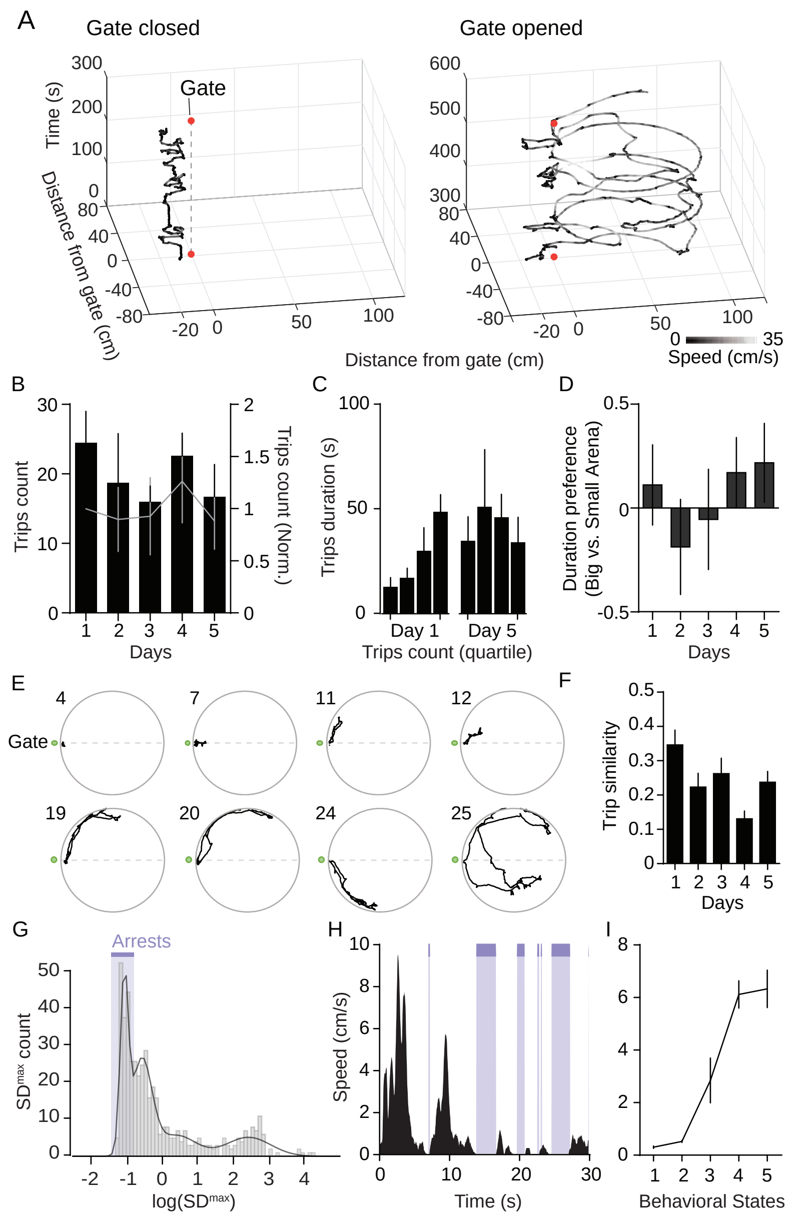
A
Gate closed
Gate opened
Gate
Time (s)
Distance from gate (cm)
Speed (cm/s)
B
Trips count
Trips count (Norm.)
Days
C
Trips duration (s)
Day 1
Day 5
Trips count (quartile)
D
Duration preference
(Big vs. Small Arena)
Days
E
Gate
F
Trip similarity
Days
G
Arrests
SDmax count
log(SDmax)
H
Speed (cm/s)
Time (s)
I
Behavioral States

Figure SI 1

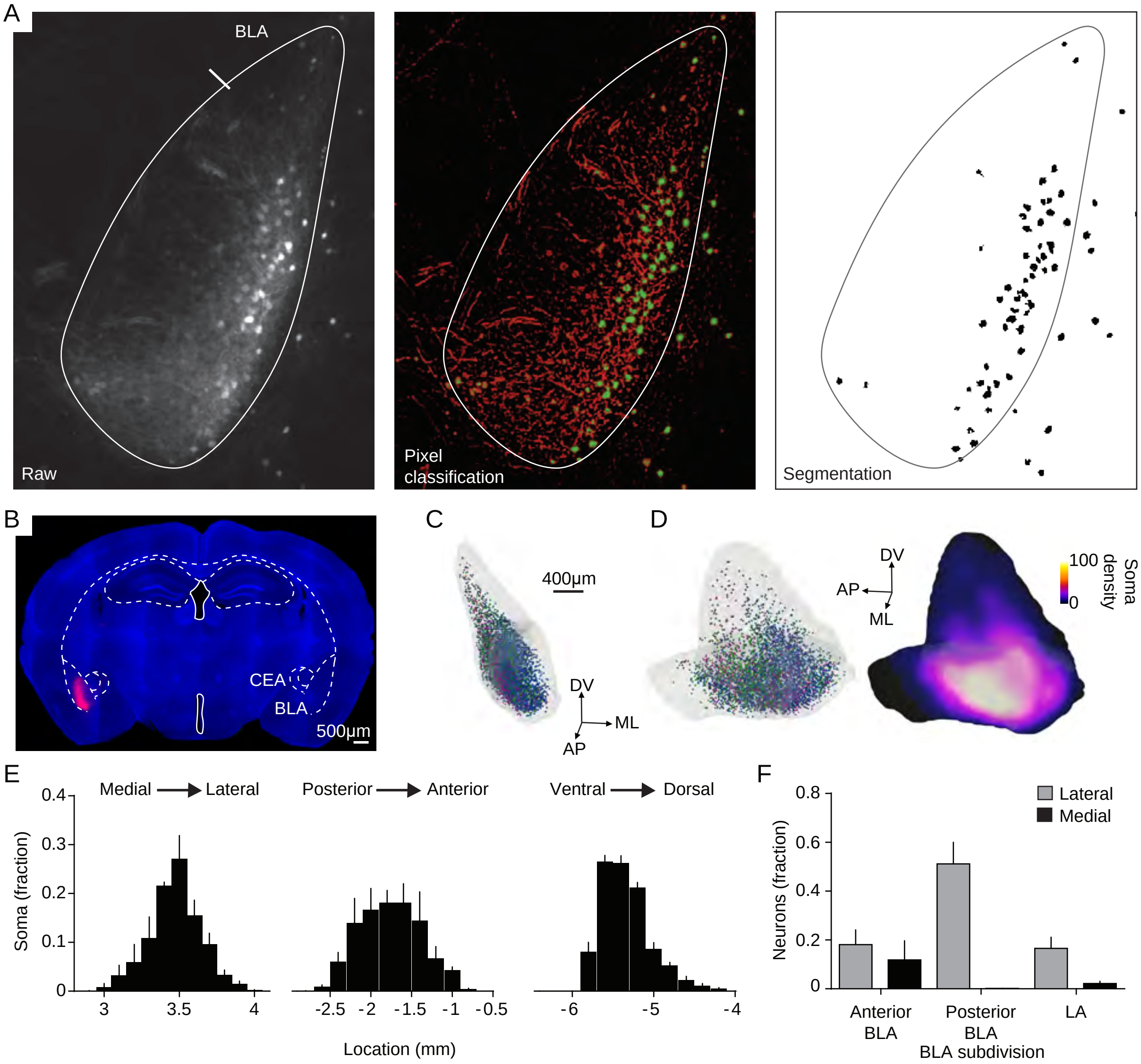
A
BLA
Raw
Pixel
classification
Segmentation
B
CEA
BLA
500µm
C
400µm
DV
ML
AP
D
DV
AP
ML
100
0
Soma
density
E
Medial → Lateral
Posterior → Anterior
Ventral → Dorsal
Soma (fraction)
0.4
0.3
0.2
0.1
0
3
3.5
4
-2.5 -2 -1.5 -1 -0.5
-6
-5
-4
Location (mm)
F
0.8
0.6
0.4
0.2
0
Neurons (fraction)
Lateral
Medial
Anterior
BLA
Posterior
BLA
LA
BLA subdivision

Figure SI 2

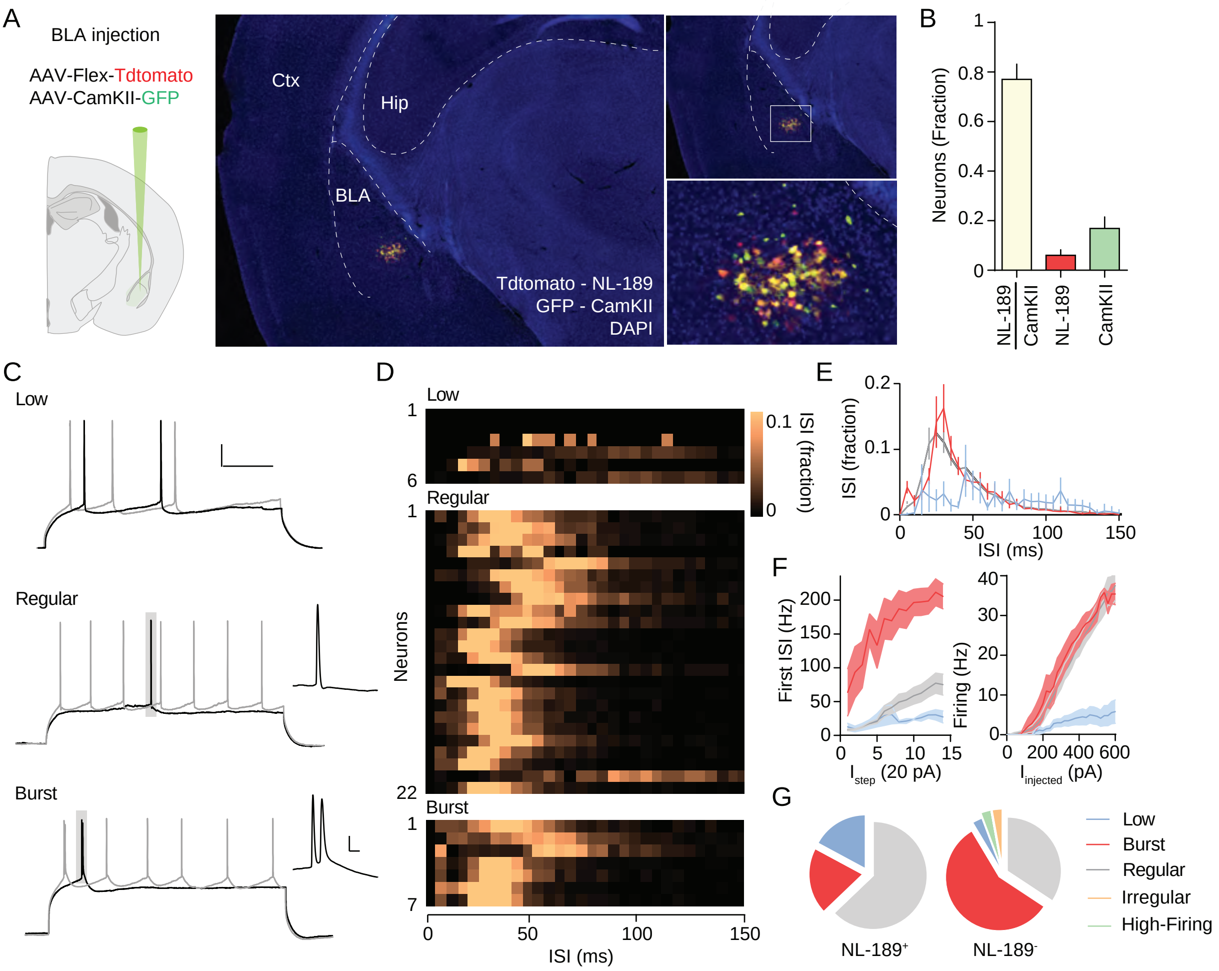

A
BLA injection
AAV-Flex-Tdtomato
AAV-CamKII-GFP
Ctx
Hip
BLA
Tdtomato - NL-189
GFP - CamKII
DAPI
B
Neurons (Fraction)
NL-189 CamKII
NL-189
CamKII
C
Low
Regular
Burst
D
Low
Regular
Burst
Neurons
ISI (fraction)
ISI (ms)
E
ISI (fraction)
ISI (ms)
F
First ISI (Hz)
Istep (20 pA)
Firing (Hz)
Iinjected (pA)
G
Low
Burst
Regular
Irregular
High-Firing
NL-189+
NL-189-


Figure SI 3

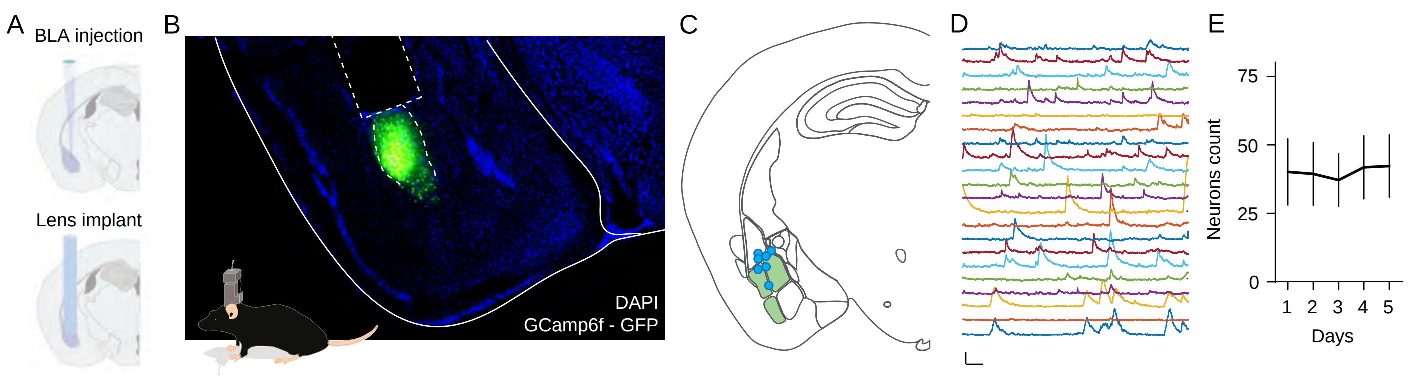
A
BLA injection
Lens implant
B
DAPI
GCamp6f - GFP
C
D
E
75
50
25
0
Neurons count
1 2 3 4 5
Days

Figure SI 4

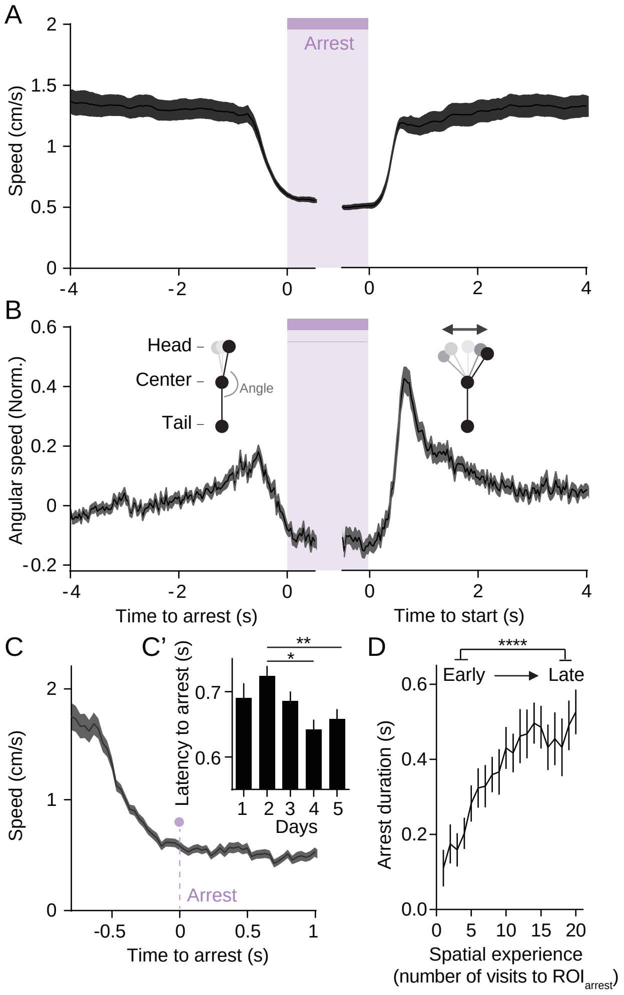
A
Arrest
Speed (cm/s)
B
Head
Center
Angle
Tail
Angular speed (Norm.)
Time to arrest (s)
Time to start (s)
C
Speed (cm/s)
Arrest
Time to arrest (s)
C’
Latency to arrest (s)
Days
D
Early
Late
Arrest duration (s)
Spatial experience
(number of visits to ROI$_{arrest}$)

Figure SI 5

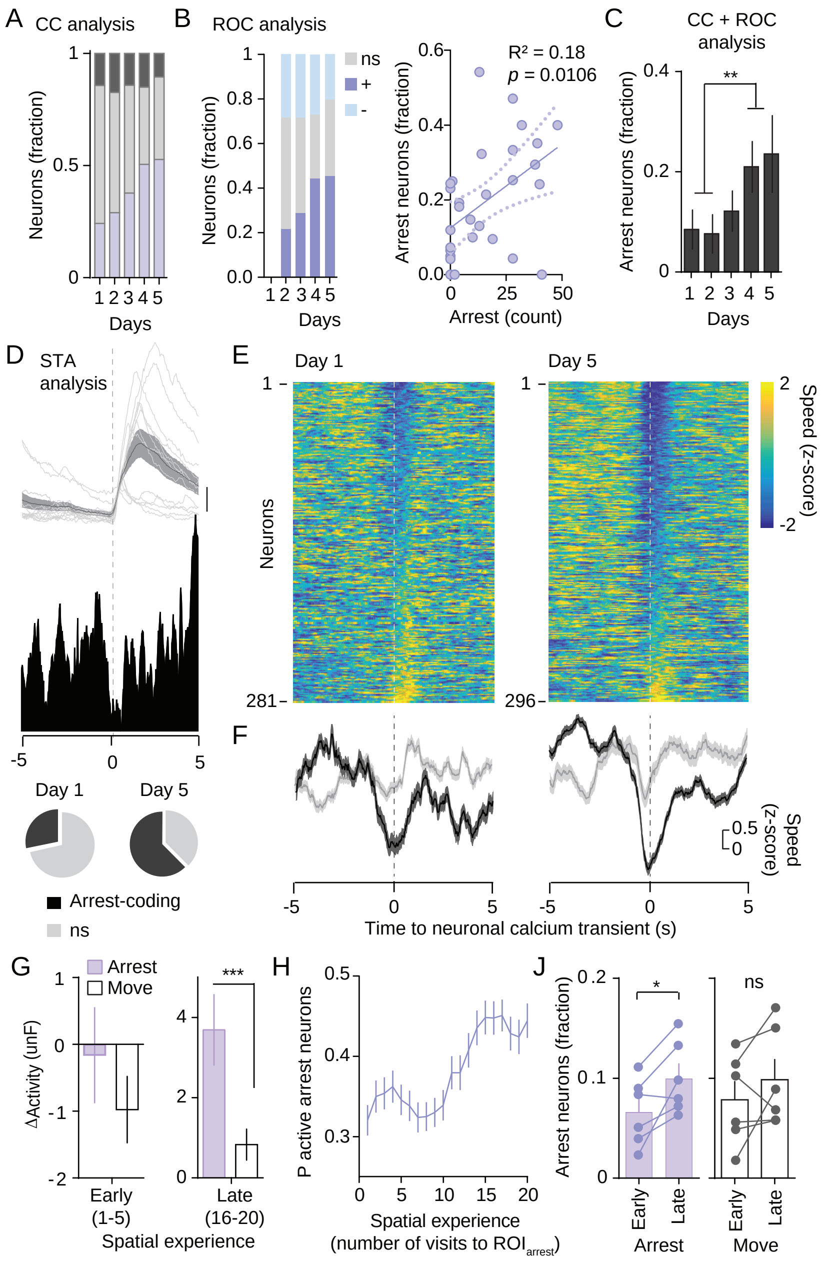

A CC analysis
B ROC analysis
C CC + ROC analysis
Neurons (fraction)
Days
ns
+
-
R² = 0.18
p = 0.0106
Arrest neurons (fraction)
Arrest (count)
**
D STA analysis
E Day 1
Day 5
Neurons
Speed (z-score)
Day 1
Day 5
Arrest-coding
ns
F
Time to neuronal calcium transient (s)
Speed (z-score)
G Arrest
Move
ΔActivity (unF)
***
Early (1-5)
Late (16-20)
Spatial experience
H P active arrest neurons
Spatial experience (number of visits to ROI arrest)
J Arrest neurons (fraction)
*
ns
Early
Late
Arrest
Move


Figure SI 6

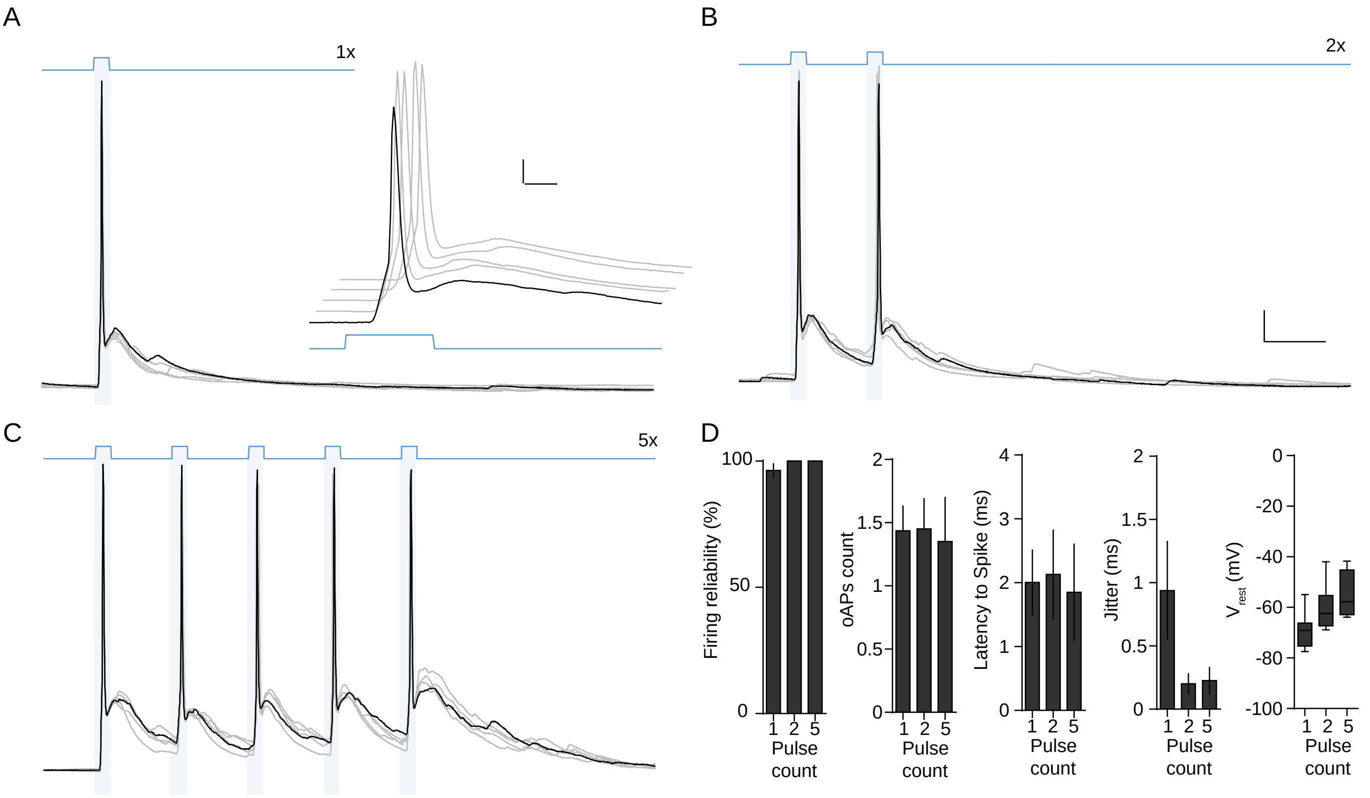

A
1x
B
2x
C
5x
D
Firing reliability (%)
100
50
0
oAPs count
2
1.5
1
0.5
0
Latency to Spike (ms)
4
3
2
1
0
Jitter (ms)
2
1.5
1
0.5
0
$V_{rest}$ (mV)
0
-20
-40
-60
-80
-100
1 2 5
Pulse count


Figure SI 7

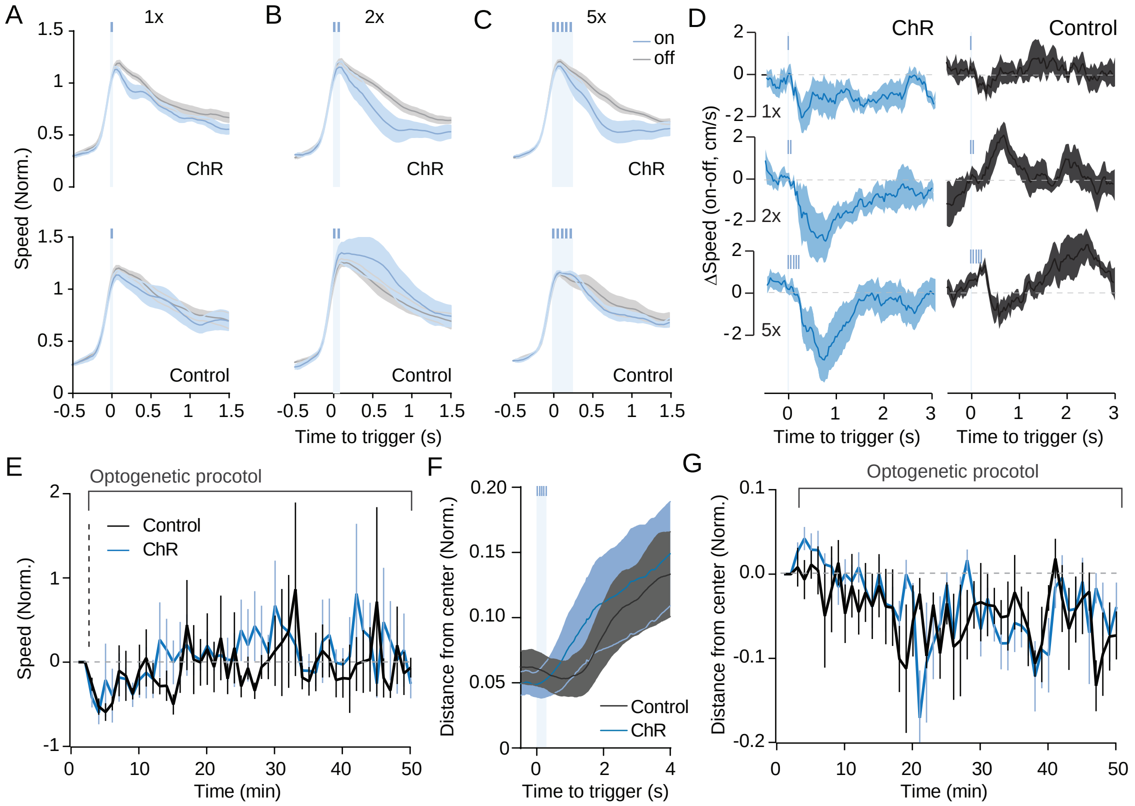

A
B
C
D
E
F
G
1x
2x
5x
on
off
ChR
Control
Speed (Norm.)
Time to trigger (s)
ΔSpeed (on-off, cm/s)
Optogenetic procotol
Time (min)
Distance from center (Norm.)


Figure SI 8

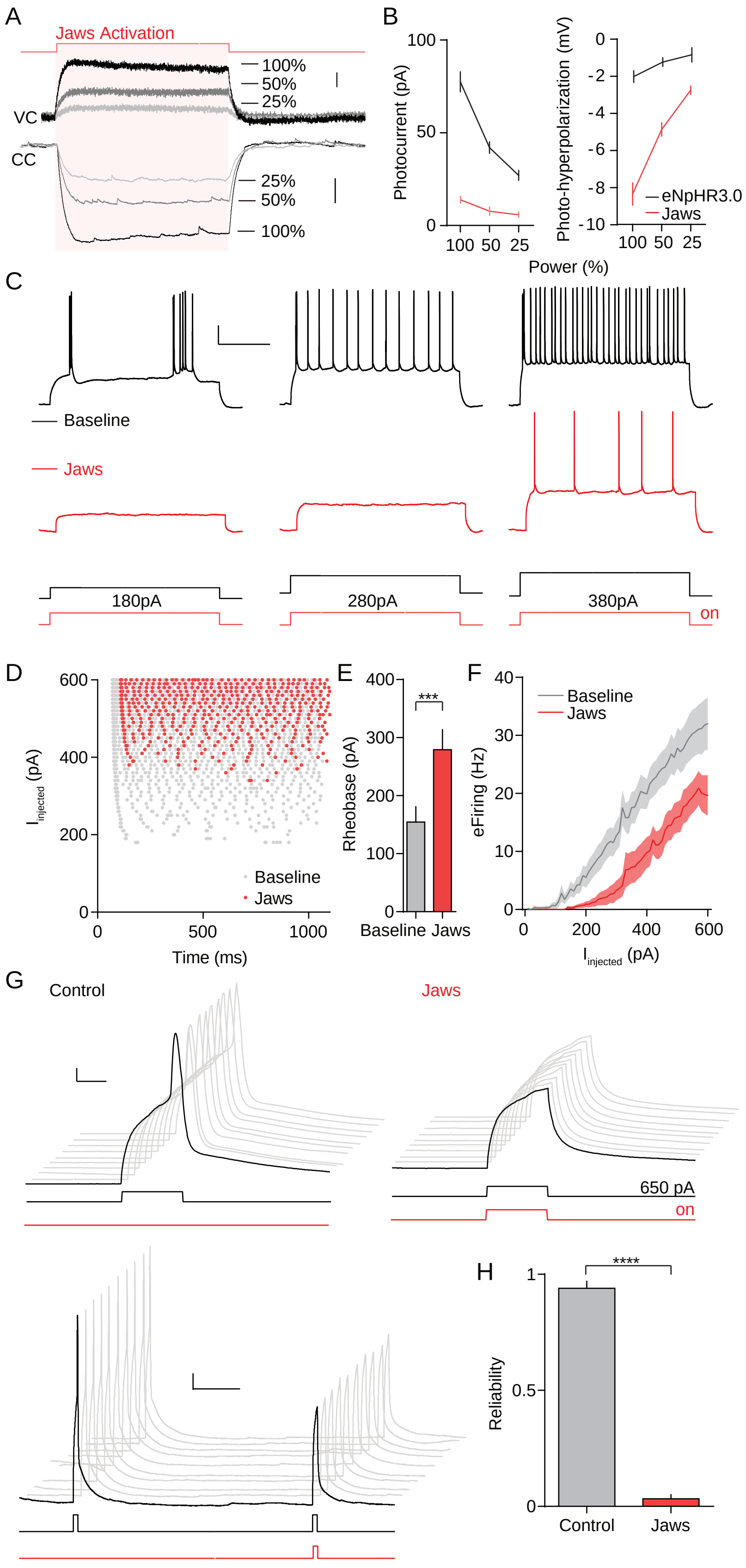
A
Jaws Activation
VC
CC
100%
50%
25%
25%
50%
100%
B
Photocurrent (pA)
Photo-hyperpolarization (mV)
eNpHR3.0
Jaws
Power (%)
C
Baseline
Jaws
180pA
280pA
380pA
on
D
Iinjected (pA)
Time (ms)
Baseline
Jaws
E
Rheobase (pA)
***
Baseline Jaws
F
eFiring (Hz)
Baseline
Jaws
Iinjected (pA)
G
Control
Jaws
650 pA
on
H
Reliability
****
Control
Jaws

Figure SI 9

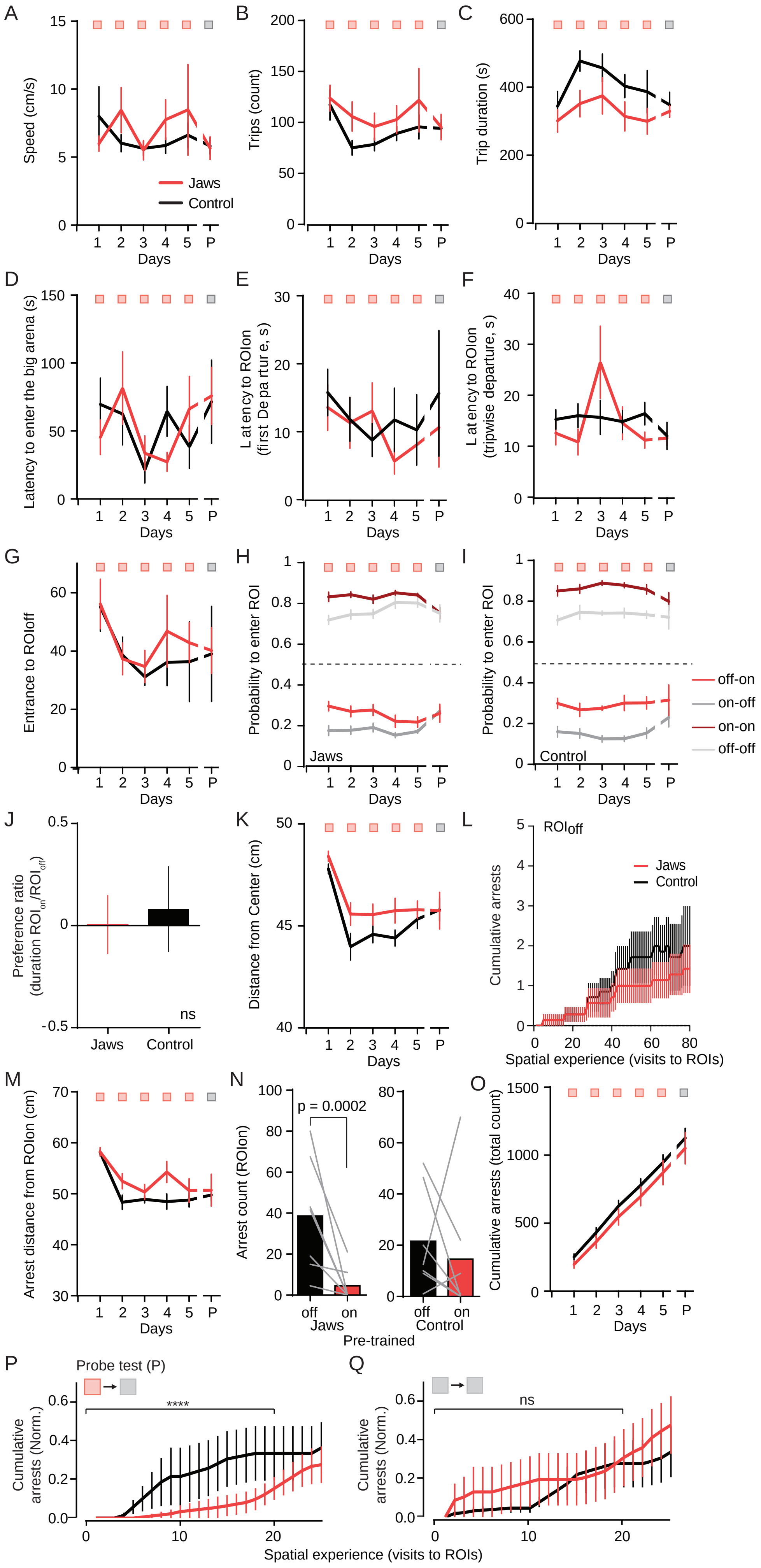
A
B
C
D
E
F
G
H
I
J
K
L
M
N
O
P
Q
Speed (cm/s)
Trips (count)
Trip duration (s)
Jaws
Control
Days
Latency to enter the big arena (s)
Latency to ROIon (first Departure, s)
Latency to ROIon (tripwise departure, s)
Entrance to ROIoff
Probability to enter ROI
off-on
on-off
on-on
off-off
Preference ratio (duration ROIon/ROIoff)
ns
Distance from Center (cm)
ROIoff
Cumulative arrests
Spatial experience (visits to ROIs)
Arrest distance from ROIon (cm)
Arrest count (ROIon)
p = 0.0002
off
on
Pre-trained
Cumulative arrests (total count)
Probe test (P)
Cumulative arrests (Norm.)
****

Figure SI 10

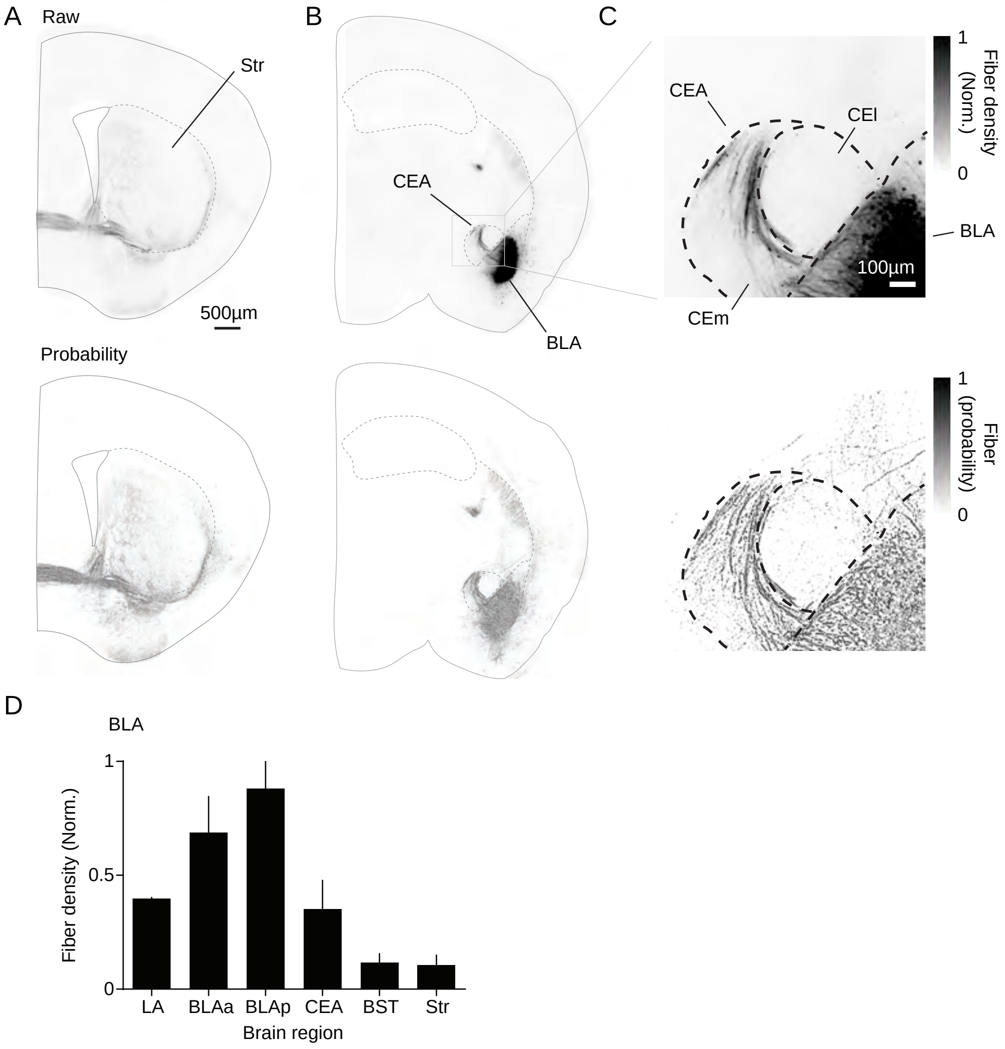
A
Raw
Str
500µm
Probability
B
CEA
BLA
C
CEA
CEl
CEm
BLA
100µm
1
Fiber density (Norm.)
0
1
Fiber (probability)
0
D
BLA
Fiber density (Norm.)
1
0.5
0
LA
BLAa
BLAp
CEA
BST
Str
Brain region

Figure SI 11

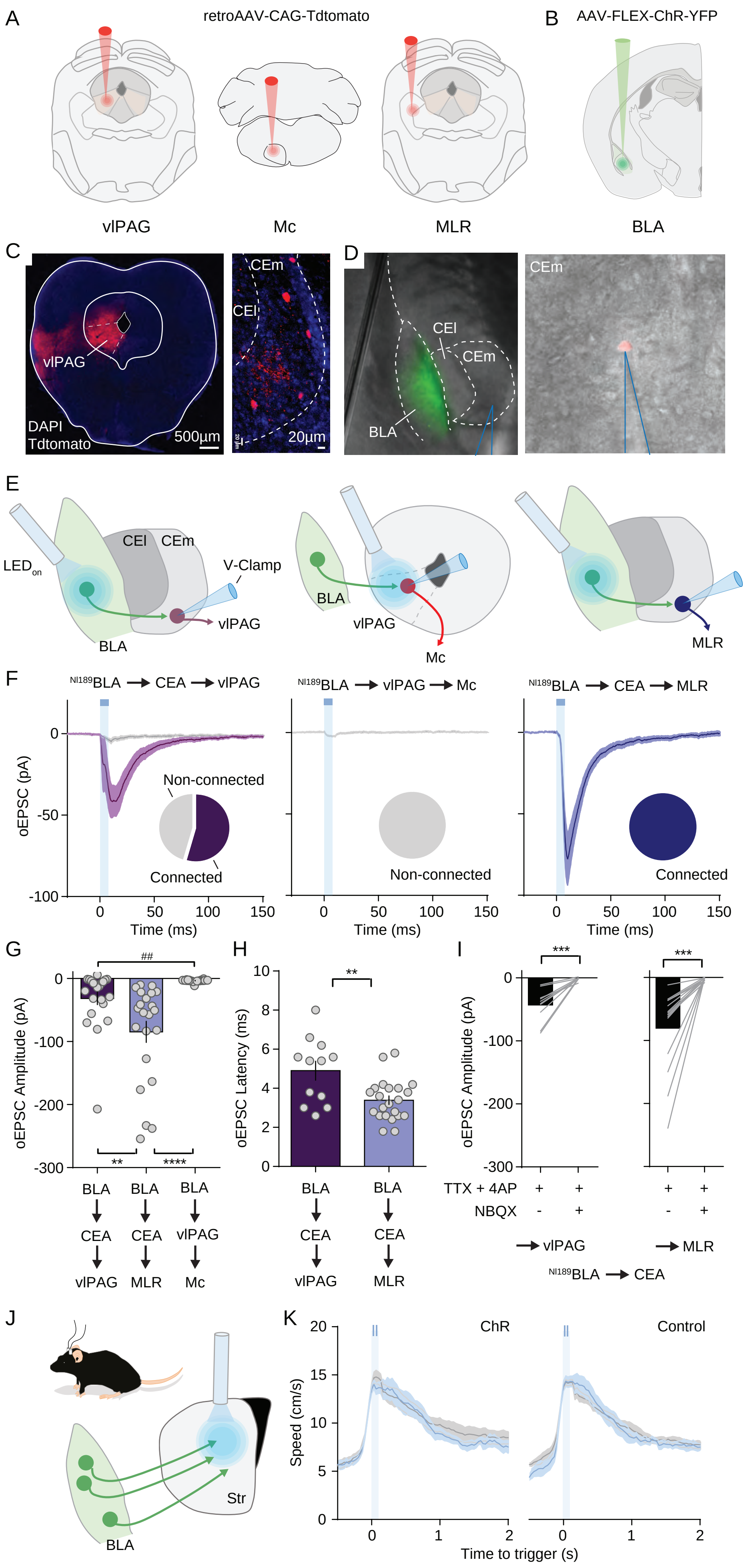
A
retroAAV-CAG-Tdtomato
B
AAV-FLEX-ChR-YFP
vlPAG
Mc
MLR
BLA
C
vlPAG
DAPI
Tdtomato
500µm
CEm
CEl
20µm
D
CEl
CEm
BLA
CEm
E
LED on
CEl
CEm
V-Clamp
vlPAG
BLA
BLA
vlPAG
Mc
MLR
F
NI189BLA → CEA → vlPAG
NI189BLA → vlPAG → Mc
NI189BLA → CEA → MLR
oEPSC (pA)
0
-50
-100
Non-connected
Connected
Non-connected
Connected
0
50
100
150
Time (ms)
G
##
oEPSC Amplitude (pA)
0
-100
-200
-300
**
****
BLA → CEA → vlPAG
BLA → CEA → MLR
BLA → vlPAG → Mc
H
oEPSC Latency (ms)
10
8
6
4
2
0
**
BLA → CEA → vlPAG
BLA → CEA → MLR
I
***
***
oEPSC Amplitude (pA)
TTX + 4AP
NBQX
→ vlPAG
→ MLR
NI189BLA → CEA
J
Str
BLA
K
ChR
Control
Speed (cm/s)
20
15
10
5
0
Time to trigger (s)

Figure SI 12

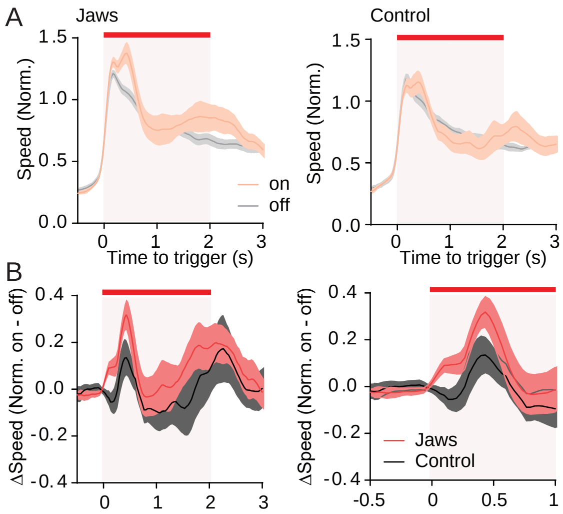

A
Jaws
Control
Speed (Norm.)
1.5
1.0
0.5
0.0
0
1
2
3
on
off
Time to trigger (s)
B
ΔSpeed (Norm. on - off)
0.4
0.2
0.0
-0.2
-0.4
-0.5
0.5
Jaws
Control


Figure SI 13

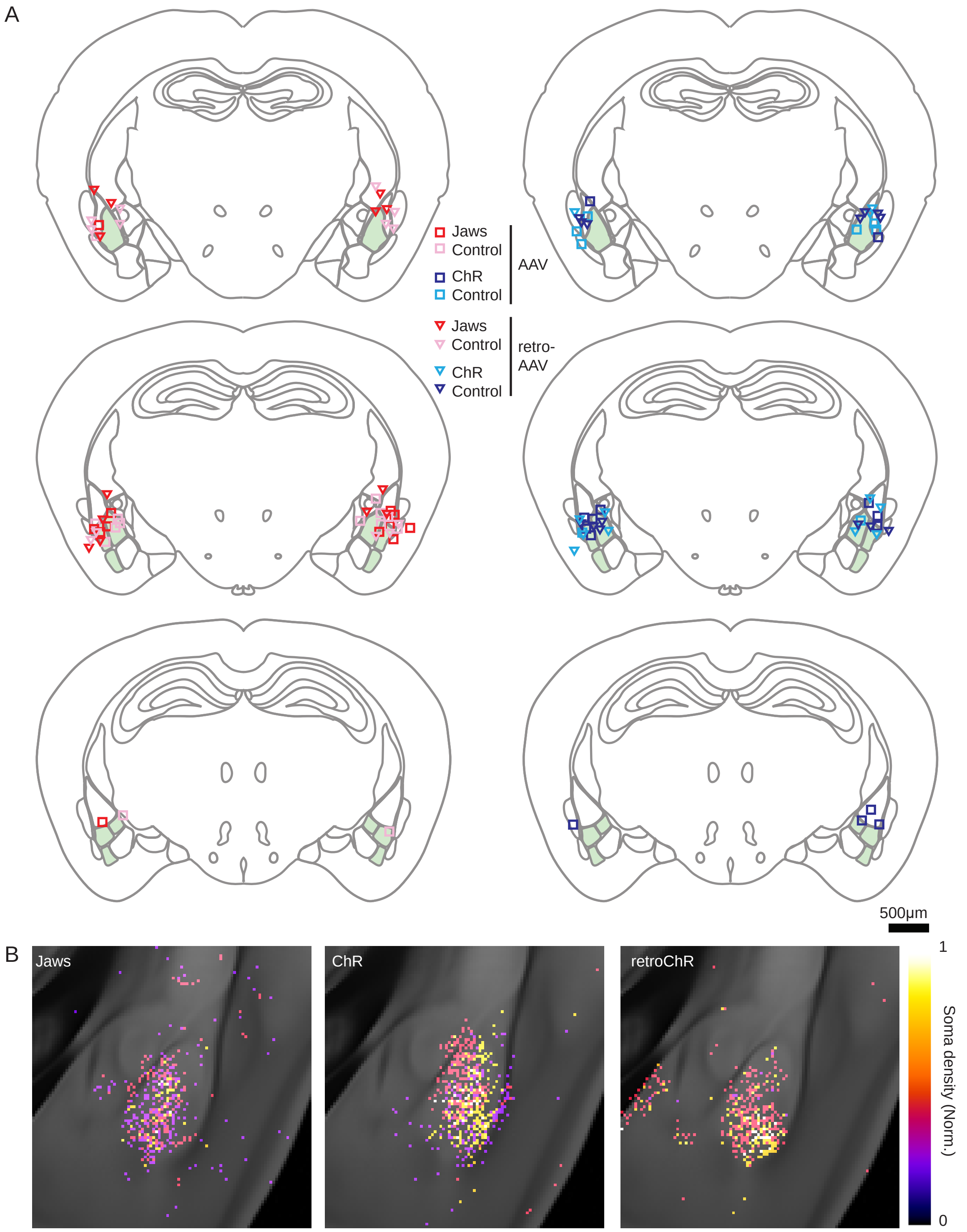
A
Jaws
Control
ChR
Control
AAV
Jaws
Control
ChR
Control
retro-
AAV
500µm
B
Jaws
ChR
retroChR
1
Soma density (Norm.)
0

Figure SI 14
